# Genomic And Molecular Characterization Of The *Wheat Streak Mosaic Virus Resistance Locus 2* (*Wsm2*) In Common Wheat (*Triticum Aestivum*. L)

**DOI:** 10.1101/2022.04.26.489503

**Authors:** Yucong Xie, Punya Nachappa, Vamsi Nalam, Stephen Pearce

**Affiliations:** Department of Soil and Crop Sciences, Colorado State University, Fort Collins, CO 80523, USA; Department of Biology, Duke University, Durham, North Carolina 27708, USA; Department of Agricultural Biology, Colorado State University, Fort Collins, CO 80523, USA; Rothamsted Research, Harpenden, Hertfordshire, AL5 2JQ, UK

**Keywords:** *Wheat streak mosaic virus*, *Wsm2*, Wheat, *RPM1*, Structural variation.

## Abstract

*Wheat streak mosaic virus* is an economically important viral pathogen that threatens global wheat production, particularly in the Great Plains region of the United States. The *Wsm2* locus confers resistance to WSMV and has been widely deployed in common wheat varieties adapted to this region. Characterizing the underlying causative genetic variant would contribute to our understanding of viral resistance mechanisms in crops and aid the development of perfect markers for breeding.

In this study, *Wsm2* flanking markers were mapped to a 4.0 Mbp region of the wheat reference genome containing 142 candidate genes. Haplotype analysis in seventeen wheat genotypes collected from different agroecological zones indicated that *Wsm2* lies in a dynamic region of the genome with extensive structural variation, and that it is likely absent from available genome assemblies of common wheat varieties.

Exome sequencing of the variety ‘Snowmass,’ which carries *Wsm2*, revealed several loss-of-function mutations and copy number variants in the 142 candidate genes within the *Wsm2* interval. Six of these genes are differentially expressed in ‘Snowmass’ compared to ‘Antero,’ a variety lacking *Wsm2*, including a gene that encodes a nucleotide-binding site leucine-rich repeat (NBS-LRR) type protein with homology to RPM1. A *de novo* assembly of unmapped RNA-seq reads identified nine transcripts expressed only in ‘Snowmass,’ three of which are also induced in response to WSMV inoculation. This study sheds light on the variation underlying *Wsm2* and provides a list of candidate genes for subsequent validation.

## 1 INTRODUCTION

Common wheat (*Triticum aestivum* L.) provides approximately 20% of the calories and proteins consumed by the human population (FAOSTAT, 2020). First observed in 1922, *Wheat streak mosaic virus* (WSMV) is the type species of the genus *Tritimovirus* within the family *Potyviridae* (Stenger et al., 1998; Singh et al., 2018) and is an economically important viral pathogen that threatens wheat production around the globe (Navia et al., 2013). In the United States, WSMV mainly affects wheat grown in the Great Plains region, causing average annual yield losses of approximately 5%, although severe localized infections can result in complete crop failure (Singh and Kundu, 2018; McKelvy et al., 2021). Once infected with WSMV, wheat leaves exhibit a characteristic yellow and green streaked mosaic pattern (Hadi et al., 2011). For winter wheat varieties, symptoms are most severe when infection occurs during tillering and include stunting, poor fertility, and reduced grain set (Hunger et al., 1992).

The transmission vector for WSMV is the eriophyid wheat curl mite (WCM) *Aceria tosichella* Keifer, which has a body length of ∼200 μm and is spread between plants by the wind (Slykhuis, 1955). WSMV can be acquired by WCM from infected host plants during a 10- to 30-minute feeding time and remains active in WCM for 7-9 days (Orlob, 1966). Upon landing on wheat plants, the WCM remains hidden in rolled and curled leaves and leaf sheaths, where it can feed and survive for several months (Navia et al., 2013). As a result, miticides are ineffective in controlling WCM populations (Navia et al., 2013). Moreover, volunteer wheat, other monocots and wild weeds, including oats (*Avena sativa*), barley (*Hordeum vulgare*), rye (*Secale cereale*), corn (*Zea mays*), and foxtail millet (*Setaria italica*), can serve as a ‘green bridge’ for WCM to complete their life cycle between wheat cropping seasons (Singh and Kundu, 2018). This broad host range makes it ineffective and impractical for many growers to use cultural practices to eradicate WCM from infected fields (Singh et al., 2018). Therefore, the most effective long-term strategy to prevent damage caused by WCM and WSMV is to develop wheat cultivars with genetic resistance to the WSMV-WCM disease complex (Harvey et al., 1999; Nachappa et al., 2021).

To date, four quantitative trait loci (QTL) associated with WCM resistance (known as *Curl mite colonization* or *Cmc* genes) have been identified from grass species and ancestors of common wheat (Thomas and Conner, 1986; Whelan and Hart, 1988; Malik et al., 2003). Although these resistance alleles inhibit WCM reproductive potential and reduce its transmission rate in the field, their effectiveness varies between WCM populations and environmental conditions (Murugan et al., 2011; Dhakal et al., 2017). Moreover, all four *Cmc* loci are derived from alien introgressions and are associated with reduced yields due to linkage drag (Harvey et al., 1999).

In addition to introgressing genetic resistance to WCM, four QTL (*Wsm1, Wsm2*, *Wsm3,* and *c2652*) for WSMV resistance have been identified (Haley et al., 2002; Sharp et al., 2002; Haber et al., 2006; Divis et al., 2006). Both *Wsm1* and *Wsm3* originated in intermediate wheatgrass (*Thinopyrum intermedium*) and were transferred into common wheat varieties through alien translocation (Wells et al., 1982; Friebe et al., 2009; Liu et al., 2011; Danilova et al., 2017). When deployed in elite varieties, these alleles confer a yield penalty due to linkage drag, limiting their value in wheat breeding programs. For example, in the absence of WSMV infection, *Wsm1* confers a yield penalty of up to 30% (Seifers et al., 1995; Sharp et al., 2002). Although *c2652* was identified from a hard red spring wheat population (Haber et al., 2006), it has not been utilized in wheat germplasm development to date. The most widely deployed QTL is *Wsm2*, which was first identified in the wheat breeding line CO960293-2 and most likely originated in a common wheat background (Haley et al., 2002).

Over the last 15 years, *Wsm2* has provided strong resistance to WSMV (Seifers et al., 2006; Lu et al., 2012), leading to low WSMV incidence in field conditions (McKelvy et al., 2021). However, WSMV resistance-breaking strains have been reported in infected wheat carrying *Wsm2* (Fellers et al., 2019; Redila et al., 2021; Albrecht et al., 2022) and from *Setaria viridis* (Kumssa et al., 2019). Although *Wsm2* is temperature sensitive and less effective in the field at temperatures above 18 (Seifers et al., 2006), there is no evidence that it has deleterious impacts on yield or other agronomic traits (Lu et al., 2012). Because of these advantages, *Wsm2* has been introduced into several common wheat varieties by recombination, including ‘RonL’ (Martin et al., 2007), ‘Snowmass’ (Haley et al., 2011), ‘Clara CL’ (Martin et al., 2014), ‘Oakley CL’ (Zhang et al., 2015), and ‘Joe’ (Zhang et al., 2016).

Linkage mapping in two F_2:3_ populations showed that the WSMV resistance conferred by *Wsm2* is controlled by a single dominant allele located on chromosome arm 3BS (Lu et al., 2011). Subsequent studies using a recombinant inbred line (RIL) population mapped the *Wsm2* locus to a 6.5 cM region (Assanga et al., 2017; Tan et al., 2017). Three SNP markers, each within 1 cM of *Wsm2*, were transformed into KASP assays and validated in a RIL population (‘CO960293’×‘TAM111’) and in two doubled haploid populations (‘RonL’ × ‘Ripper’ and ‘Snowmass’ × ‘Antero’), from which haplotypes associated with WSMV resistance and susceptibility were identified (Tan et al., 2017). A genome-wide association study (GWAS) on 597 wheat breeding lines identified ten other significant SNP markers associated with WSMV resistance (Dhakal et al., 2018). These ten SNPs mapped to a 17.1 – 18.9 Mbp telomeric region on chromosome 3B in the IWGSC RefSeq v1.0 wheat reference genome assembly, coinciding with the *Wsm2* locus. This region contains a cluster of fourteen genes encoding Bowman-Birk inhibitors (BBI) and is highly diverse between wheat varieties, with evidence of structural variation (Xie et al., 2021).

Despite the importance of *Wsm2*, the underlying causative gene has not been cloned, which will be required to develop perfect markers for breeding programs, and to further our understanding of viral resistance mechanisms in crops. In the current study, genome assemblies for the landrace ‘Chinese Spring’ (IWGSC et al., 2018; Zhu et al., 2021) and sixteen other wheat varieties were used to characterize haplotypic variation at the *Wsm2* locus and to study their association with WSMV resistance. Exome and transcriptome sequencing in the variety ‘Snowmass’, which carries *Wsm2*, identified several possible causative genes underlying this locus.

## 2 MATERIALS AND METHODS

### 2.1 Plant materials

Seeds of the wheat varieties ‘Jagger’, ‘SY Mattis’, ‘Robigus’, ‘Mace’, ‘Paragon’, ‘Landmark’, ‘Stanley’, ‘Claire’, ‘Weebill’, ‘Cadenza’, ‘Kronos’, and ‘Chinese Spring’ were obtained from Seedstor (https://www.seedstor.ac.uk) and used to perform WSMV phenotyping assays. A *Triticum aestivum* L. doubled haploid (DH) population (n = 116) produced by Heartland Plant Innovations Inc. was developed by wheat-maize wide hybridization (Santra et al., 2017) from the parents ‘Snowmass’ (WSMV resistant) and ‘Antero’ (WSMV susceptible) and used for linkage mapping. ‘Snowmass’ and ‘Antero’ leaf tissues were used to quantify WSMV coat protein transcript levels by qRT-PCR in a time course from 0, 5, 10, and 15-days post inoculation (dpi). For the RNA-seq study, eight individuals homozygous for the *Wsm2* locus were selected from the DH population. Using GBS markers mapped to IWGSC RefSeq v1.0 genome, four individuals were shown to have an identical haplotype to ‘Snowmass’ (*Wsm2*+), and another four had an identical haplotype to ‘Antero’ (*Wsm2*-) (Table S1). Phenotype scores confirmed that *Wsm2+* DH lines are resistant to WSMV, whereas *Wsm2-*DH lines are susceptible (Table S1). Exome reads of ‘Snowmass’, ‘Antero’, ‘Brawl’, ‘Byrd’, ‘Hatcher’, ‘CO940610’, and ‘Platte’ were captured using the NimbleGen SeqCap EZ wheat whole-genome assay and sequenced as described in Jordan et al., 2015 and He et al., 2019.

### 2.2 WSMV inoculation and phenotype evaluation

An isolate of WSMV originally collected from Akron, Colorado, in 2017 was propagated in the greenhouse by mechanically inoculating the susceptible winter wheat genotype ‘Longhorn’ every six months. Leaf tissues with a yellow streaking or mosaic pattern typical of WSMV were collected, frozen at -80, and used to prepare fresh inoculum. The inoculum was prepared with 1:10 (w/v) dilution of the WSMV-infected wheat leaf tissue and 0.01 M potassium phosphate buffer (pH 7.4) and inoculated on two-week-old seedlings. WSMV phenotyping was performed on two-week-old wheat plants grown in a PGR15 growth chamber (Conviron, Manitoba, Canada) in a 12 h photoperiod with temperatures set to 18 day/15 night. Mechanical inoculation was performed using a soft sponge soaked with inoculum that was gently rubbed on the surface of wheat seedling leaves that were previously dusted with carborundum powder. Mock inoculation with phosphate buffer was used as a control. Phenotyping was evaluated two and three weeks after inoculation of WSMV by examining visual symptoms based on a 1-5 scale; (1 = no chlorosis; 2 = a few chlorotic streaks; 3 = moderate mosaic; 4 = severe mosaic; 5 = severe mosaic, necrosis, and yellowing) (Tan et al., 2017). Genotypes with mean scores 2 were considered resistant, and plants with scores > 2 were considered susceptible.

### 2.3 Haplotype analysis for within-species variation at the *Wsm2* locus

The *Wsm2* flanking markers wsnp_Ex_c3005_5548573 (IWA3260) and BS00026471_51 (IWB7629), one left boundary flanking marker Ku_c663_1869), and three other markers tightly linked to *Wsm2* (IAAV6442, BS00018764_51, and IWA7647, Table S2) were used for haplotype analysis. BLAST alignment of 100 bp surrounding sequences (Total 201 bp sequence with the SNP at position 101 bp) of these markers was used to identify the physical position of *Wsm2* on the wheat reference genomes IWGSC RefSeq v1.0 (IWGSC 2018) and RefSeq v2.1 (Zhu et al., 2021). These SNPs were also mapped to the genome assemblies of sixteen other wheat varieties (Mace, Lancer, CDC Stanley, CDC Landmark, Julius, Norin61, ArinaLrFor, Jagger, Cadenza, Paragon, Kronos, Robigus, Claire, Spelt, Weebill, SY Mattis) to analyze genomic variation using the Galaxy platform (Afgan et al., 2018).

For validation of the genomic variation between varieties, the genomic sequence of *Wsm2* in these wheat cultivars were extracted using the ‘bedtools’ getfasta command. The FASTA files of *Wsm2* in different wheat varieties were subjected to pairwise alignment and dot plots were generated with ‘D-genies’ using the minimap function (Cabanettes and Klopp, 2018).

### 2.4 Linkage mapping analysis for the DH population

The DH population was subjected to genotyping-by-sequencing (GBS, Elshire et al., 2011), and data were processed as described in Liu et al., 2016. The GBS markers were mapped to the wheat reference genome IWGSC RefSeq v1.0 (IWGSC 2018) and annotated for their position. For example, marker S3B_16589830 is at position 16,589,830 bp on wheat chromosome 3B. Quantitative trait loci (QTL) analysis was performed with the R version 4.0.3 packages ‘R/qtl’ (Arends et al., 2010) and ‘ASMap’ (Taylor and Butler, 2017) using the mean phenotyping scores from four biological replicates of each DH line.

### 2.5 Exome capture analysis for genetic variation underlying *Wsm2* locus

The 150 bp paired-end Illumina reads were filtered for quality using ‘fastp’ (Chen et al., 2018). Reads were then aligned to the wheat reference IWGSC RefSeq v1.0 assembly using ‘bowtie2’ v. 2.3.5 (Langmead and Salzberg, 2012) with the following parameters: -k 2 -N 1 -L 22 -D 20 -R 3. The alignments were subjected to ‘samtools’ v1.11 to generate sorted BAM files and then ‘bcftools’ v1.11 (Danecek et al., 2021) was used to call variants within *Wsm2* locus with the ‘mpileup’ command. The ‘SnpEff’ (Cingolani et al., 2012) tool was used to predict the effects of genetic variants, including SNPs, indels, and multiple-nucleotide polymorphisms. The sorted BAM files from ‘Antero’, ‘Brawl’, ‘Byrd’, ‘Hatcher’, ‘CO940610’, and ‘Platte’ were used as a reference set to assay copy number variation (CNV) for the test sample ‘Snowmass’ (*Wsm2*+) using the ‘ExomeDepth R’ package v.1.1.12 (Plagnol et al., 2012). The default parameters were used, except for transition probability = 0.001 (“CallCNVs”) and min.overlap = 0.01 (“AnnotateExtra”).

To calculate the total variant number in defined regions, the sorted BAM files were subjected to ‘bcftools’ to generate VCF files and specify the defined region using (-r) in the ‘bcftools mpileup’ command. The output VCF files were then subjected to ‘SnpEff’ to predict the total number of variants within each region. The total exome or protein coding gene length was calculated using GFF3 file from IWGSC RefSeq v1.1 (http://ftp.ensemblgenomes.org/pub/plants/release-52/gff3/triticum_aestivum/) to extract the start and end position for each ‘gene’ feature and sum the length of each ‘gene’ feature within the selected interval. The variant rate was calculated by dividing total exome length by total variant number (one variant / bp exome length). The total variant rate for ‘Snowmass’ was calculated for each individual chromosome and for the 4.0 Mbp *Wsm2* interval. In addition, six other selected 4.0 Mb intervals on telemetric or centromeric region of chromosome 3B as well as the homologous region of *Wsm2* on three other chromosomes (3B:40-44Mb, 3B:200-204Mb, 3B:345-349Mb,1A:15-19Mb, 2B:15-19Mb, and 6D:15-19Mb), together with the *Wsm2* interval in six other wheat varieties were used as comparisons for the overall variant rate.

### 2.6 WSMV quantification

Two weeks after germination, ‘Antero’ (*Wsm2*-, WSMV susceptible) and ‘Snowmass’ (*Wsm2*+, WSMV resistant) plants were subjected to WSMV inoculation as described above. Leaf samples were taken by cutting the whole leaf tissues from five biological replicates over the time course of 0, 5, 10, and 15 dpi. Leaf tissues were ground and homogenized in liquid nitrogen for subsequent total RNA isolation with the Spectrum^TM^ Plant Total RNA Isolation Kit (Sigma, USA), followed by on-column DNase I digestion treatment (Sigma-Aldrich) to remove genomic DNA according to the manufacturer’s instructions. The one-step qRT-PCR reaction was carried out using the TaqMan® RNA-to-Ct^TM^ 1-step kit (Applied Biosystems^TM^) on a QuantStudio^TM^3 Real-Time PCR system (Applied Biosystems). Reaction conditions and parameters were described previously (Price et al., 2010). WSMV was detected via qRT-PCR using coat protein specific primers and the probe listed in Table S3. To quantify the absolute amount of WSMV coat protein transcript levels, a regression line was plotted using plant RNA that contains 10 ng/µL of WSMV and subjected to 10-fold serial dilutions to 1 × 10^-5^ ng/µL. The C_T_ values for each dilution were plotted against total RNA transcript levels, and the regression line was considered at R^2^ > 0.996.

### 2.7 RNA-seq library preparation

Sixteen samples were collected for the RNA-seq experiment, including four biological replicates of two genotypes (*Wsm2*+ and *Wsm2*-) and two treatments (mock inoculation with phosphate buffer (C) and WSMV inoculation with infected tissue (T)). RNA-seq samples were labeled as *Wsm2*- (C), *Wsm2*- (T), *Wsm2*+ (C) and *Wsm2*+ (T). Whole leaf tissue was harvested at 10 dpi, stored at -80 □, and ground to a homogenized fine powder in liquid nitrogen. Total RNA was isolated with Spectrum^TM^ Plant Total RNA Isolation Kit (Sigma, USA) and quantified using Qubit. The Agilent 2100 bioanalyzer (RNA Nano Chip, Agilent, CA) was used to check RNA integrity. The library construction and sequencing via Illumina HiSeq 2000 were performed by Novogene Co., Ltd (Sacramento, CA, USA), and approximately 150 bp paired-end raw reads were generated. Raw sequencing data is available from the NCBI Gene Expression Omnibus under accession number GSE190382.

### 2.8 Transcript abundance, differential expression (DE) and GO enrichment analysis

To quantify WSMV reads in the RNA-seq samples, the coding sequence from WSMV isolate KSHm2014 (9,384 bp) was retrieved from the NCBI database (MK318278.1, https://www.ncbi.nlm.nih.gov/). This sequence was concatenated to the IWGSC RefSeq v1.0 wheat genome (IWGSC 2018) as an additional FASTA entry, and used as the reference genome to build index files for alignments. Raw reads of each paired-end library were examined for sequence quality and adaptor sequences were removed using ‘fastp’ with default settings (Chen et al., 2018). Trimmed paired-end RNA-seq reads were aligned to the reference genome using ‘STAR’ 2.7.3 (Dobin and Gingeras, 2015) with parameters “-outFilterMismatchNmax 6 - alignIntronMax 10000”. Non-normalized reads were counted with ‘featureCounts’ (Liao et al., 2014) with parameters “-t gene -p” and used as input for the R package ‘DEseq2’ v3.14 (Love et al., 2014). Read counts of WSMV in each sample were normalized to ‘fastp’ trimmed reads of the corresponding samples for count per million (cpm) of WSMV, and log transformed into LogCPM. Differentially expressed genes (DEGs) were identified from pairwise comparisons for treatment effect (WSMV-treated *vs.* mock-treated), and for genotypic effect under each condition: Resistant *vs.* Susceptible under WSMV-treated condition 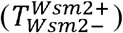 and under mock-treated condition 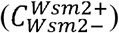. The *P*-value threshold was determined using Benjamini and Hochberg’s approach (Benjamini and Hochberg, 1995) for controlling the false discovery rate (FDR < 0.01) without controlling the log_2_ fold change (FC). Venn diagrams were drawn using VENNY software (Oliveros, 2007). Gene ontology (GO) enrichment analyses were performed with the ‘TopGO’ R package v3.14 (Alexa A, 2021) and Fisher tests were conducted to identify significant GO terms (*P* < 0.01).

### 2.9 *De novo* transcriptome assembly of unmapped reads in *Wsm2+* and presence/absence analysis

During the STAR alignment, reads that did not map to IWGSC RefSeq v1.0 were collected with parameter “-outReadsUnmapped” and assembled with ‘Trinity’ tool v2.14.0 (Grabherr et al., 2011) using the following parameters: “-seqType fq –samples_file<INPUT_FILE> -max_memory 10G -CPU 20”. The proportion of reads mapped to the assembly was assessed with ‘Bowtie2’ v2.3.5 (Langmead and Salzberg, 2012). Then ‘CD-HIT’ (cd-hit-est – c 0.95) was used to remove redundant transcripts. The assembled transcripts were used as BLASTx queries against the NCBI NR database of non-redundant proteins (cutoff: 1e-5) to annotate gene function. DEG analysis were performed on unmapped reads against assembled transcriptomes to identify differentially expressed transcripts (DETs) in four pairwise comparisons: 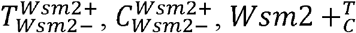, and 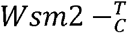.

Presence and absence (PAV) analysis was performed by BLASTn (default: 1e-3) the assembled transcript sequences against the coding sequences (CDS) of other wheat varieties using the Galaxy platform (Afgan et al., 2018). The transcript was determined to be present (+) when the top BLAST hit exhibited a CDS similarity percentage > 96%, otherwise the transcript was determined to be absent from that genome assembly. For unique transcripts in *Wsm2+* that are also present in other wheat varieties, the corresponding gene ID and physical position were extracted using the ‘Galaxy’ platform (Afgan et al., 2018).

### 2.10 Gene expression validation with qRT-PCR

Transcript levels of selected candidate DEGs identified from RNA-seq experiments were validated with qRT-PCR in the same samples used for WSMV quantification. First-strand cDNA was synthesized from 2 µg of total RNA using SuperScript IV Reverse Transcriptase Kit (Thermo Fisher Scientific, USA) according to the manufacturer’s instructions. The qRT-PCR reactions were performed using PowerUp SYBR Green Master Mix (Thermo Fisher Scientific, USA) in a 20 µL reaction with 100 ng cDNA and 1 μL of a 10 μM solution for each primer. Relative gene expression analysis was calculated using *ACTIN* as the internal control gene and 2 ^-Δ*CT*^ method was used for relative quantification. Primer efficiency and specificity were determined by analyzing amplification in a four-fold dilution series and checking the dissociation curve for a single amplified product and calculated as: Efficiency 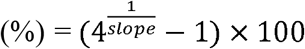. All primers in this study had an efficiency greater than 90% and are listed in Table S3.

## 3 RESULTS

### 3.1 Genomic characterization of *Wsm2* in common wheat

#### 3.1.1 Analysis of within-species genomic variation for *Wsm2* in seventeen wheat varieties

The *Wsm2* flanking markers wsnp_ex_c3005_5548573 (referred to as SNP1 hereafter) and BS00026471_51 (SNP6) were used to define the physical position of the *Wsm2* locus in genome assemblies of the landrace ‘Chinese Spring’ (Table S2). SNP1 mapped to two locations on chromosome arm 3BS, 14,985,292 bp and 26,650,801 bp. At both positions, 100 bp flanking sequences were identical except that the former carries the ‘C’ allele type, whereas the latter carries the “T” allele type for this SNP. Because SNP1 mapped to two locations, it is an ambiguous marker to define the left boundary of *Wsm2*. Instead, an alternative marker Ku_c663_1896 (SNP2) which is 0.3 cM from SNP1 in the RIL population and 680 bp downstream of SNP1 mapped uniquely at position 14,985,972 bp (Table S2) and was used to define the left boundary of *Wsm2* in subsequent analyses.

The markers SNP2 and SNP6 span a 4.0 Mbp interval on chromosome arm 3BS in both the RefSeq v1.0 (15.0 - 19.0 Mbp) and RefSeq v 2.1 (20.5 Mbp - 24.5 Mbp) genome assemblies (Table S2). In both assemblies, this region included the same 142 annotated gene models (70 high confidence, 72 low confidence, Table S4). Three other SNP markers, each within 1 cM of *Wsm2* (Tan et al., 2017), referred to as SNP3, SNP4, and SNP5, together with the flanking marker SNP6, were used to define the haplotype across the *Wsm2* region. ‘Chinese Spring’ carries the WSMV-susceptibility haplotype ‘CGTG’ (Figure 1), consistent with its mean phenotypic score of 4.0 (based on a 1-5 scale of visual symptoms where ≤ 2.0 indicates resistance, and > 2.0 indicates susceptibility) (Table 1). These results indicate that the IWGSC RefSeq v1.0 genome assembly likely does not contain the *Wsm2* genetic variant.

**Figure 1.**
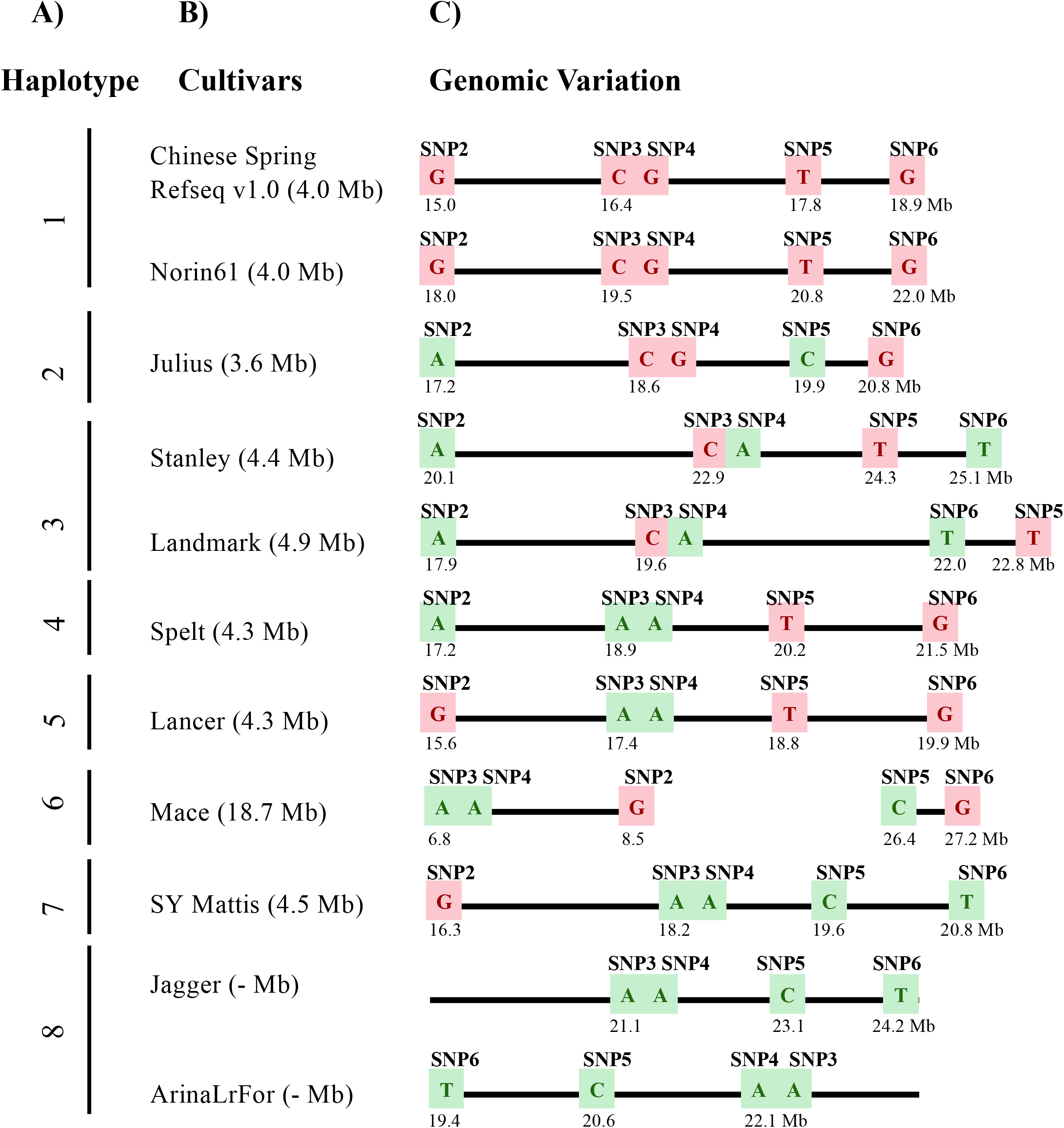
Genomic variation underlying *Wsm2* in eleven wheat varieties. **A)** Haplotypes were grouped based on allele type across five SNP markers (SNP2-SNP6). **B)** Variety names and the *Wsm2* interval size. **C)** Haplotype and genomic position based on SNP2-SNP6. The relative physical position of markers were drawn to scale (1 cm = 1 Mbp). WSMV resistant haplotype is highlighted in green and WSMV susceptible haplotypes in pink. Values below each SNP indicate the physical position (Mbp) in each respective genome assembly. The region in ‘Mace’ is not drawn to scale because of the large insertion between SNP2 and SNP5. Due to a deletion near SNP2 in ‘Jagger’ and ‘ArinaLrFor’, the *Wsm2* interval in these two varieties cannot be defined and is indicated with a dash.

**Table 1.**
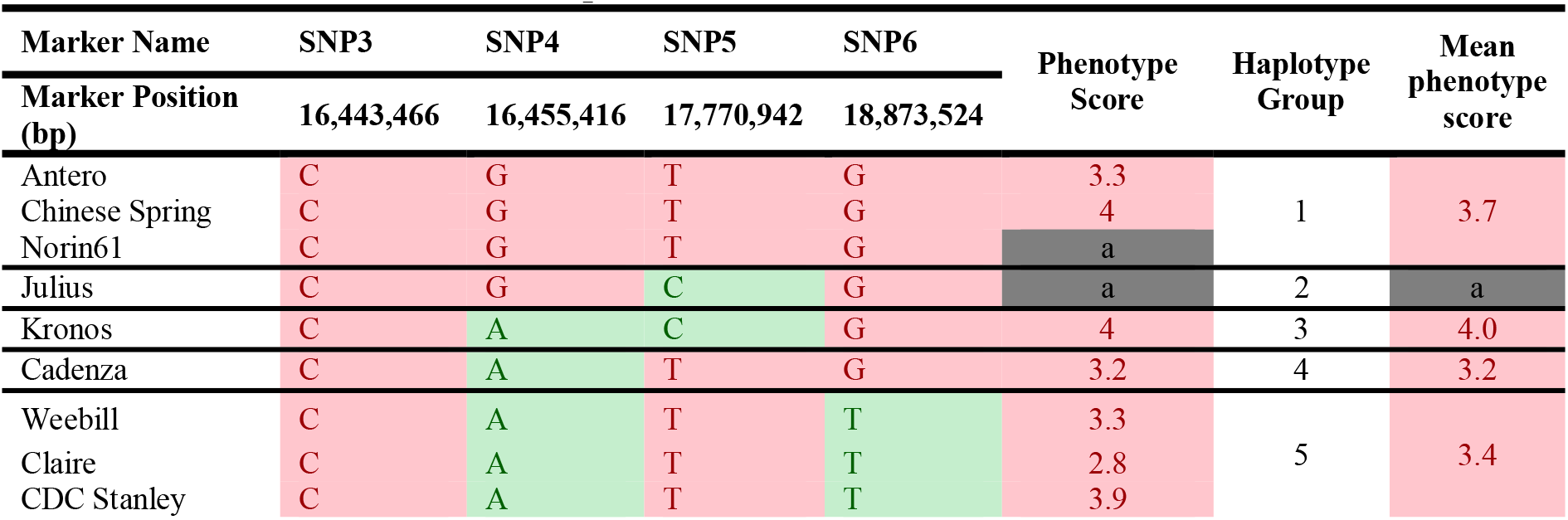
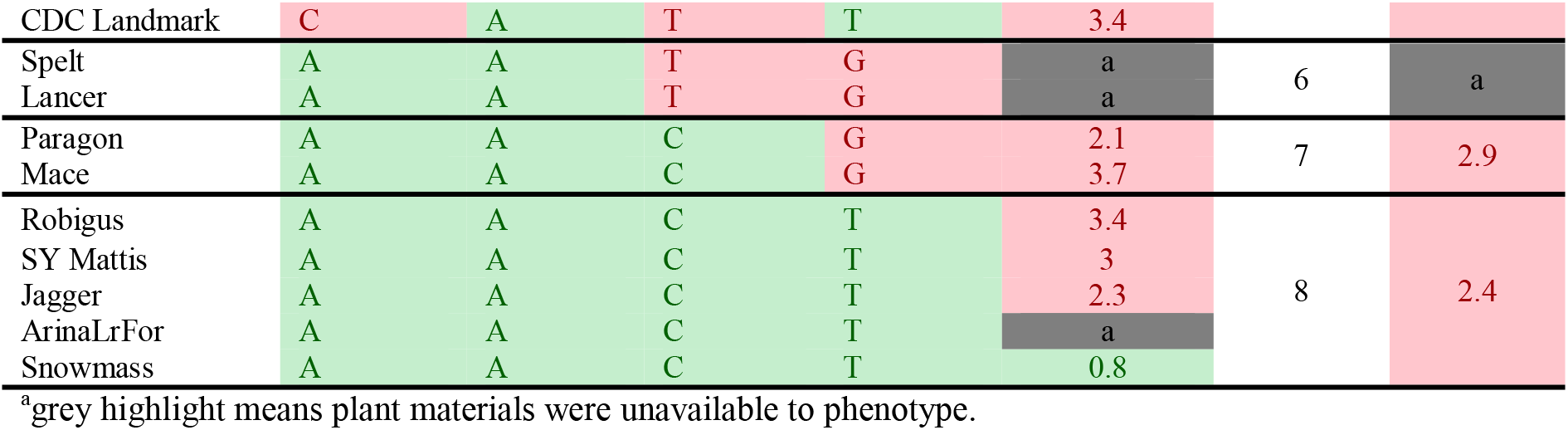
Haplotype analysis for *Wsm2* in nineteen wheat varieties. The resistant haplotype alleles or phenotype are highlighted in green, whereas susceptible haplotype alleles or phenotype are highlighted in pink. Phenotyping was performed two weeks after virus inoculation based on a 1-5 scale of visual symptoms. A mean score ≤ 2 was considered as resistant, whereas a mean score > 2 was considered as susceptible.

To compare within-species genomic diversity at the *Wsm2* locus, the corresponding genomic region was identified in ten additional wheat varieties with pseudomolecule genome assemblies by mapping the physical position of each SNP (Figure 1, Table S5). This region contained structural variation between varieties. For example, the physical distance between these five markers ranges from 3.6 Mbp in ‘Julius’ to 18.7 Mbp in ‘Mace’, due to a 17.9 Mbp insertion and inversion between markers SNP2 and SNP5 in the latter variety (Figure 1). An inversion between markers SNP3 and SNP6 was also detected in ‘ArinaLrFor’, while SNP2 is absent in ‘Jagger’ and ‘ArinaLrFor’ (Figure 1). An alignment of genomic sequence flanking this region suggests that these two varieties carry a deletion near the left boundary of *Wsm2* (Figure S1).

To further analyze *Wsm2* haplotypes, six other wheat varieties with scaffold-level genome assemblies and two winter wheat parental lines (‘Antero’ and ‘Snowmass’) were included in the analysis. Eight haplotypes in this region were identified among all nineteen wheat varieties (Table 1). The ‘CGTG’ haplotype associated with WSMV susceptibility in ‘Chinese Spring’ was also identified in ‘Norin61’ and ‘Antero’ (Table 1). Varieties with this haplotype exhibited a mean phenotypic score of 3.7 two weeks after inoculation, consistent with the association between this haplotype and WSMV susceptibility (Table 1). However, the five varieties carrying the resistant haplotype ‘AACT’ exhibited phenotypic scores ranging from 0.8 in ‘Snowmass’ to 3.4 in ‘Robigus’, with a mean score of 2.4 (Table 1), indicating that this haplotype is not consistently associated with WSMV resistance. Six other haplotypes were identified in the remaining eleven wheat varieties, all of which were susceptible to WSMV infection (mean score = 3.4, Table 1). Taken together, these results show that *Wsm2* lies in a dynamic region of the wheat genome and is likely absent from all wheat varieties with assembled genomes.

#### 3.1.2 Linkage mapping confirmed that *Wsm2* confers resistance to WSMV in ‘Snowmass’

To validate the association between *Wsm2* and WSMV resistance, linkage mapping was performed in a doubled-haploid mapping population (n = 116) derived from ‘Snowmass’ (mean phenotypic score 0.8, ‘AACT’ resistant haplotype) and ‘Antero’ (mean phenotypic score 3.3, ‘CGTG’ susceptible haplotype). Four significant QTL for WSMV resistance were identified (LOD > 3, *P* < 0.001) on chromosomes 3B, 3D, 5B and 7B (Figure 2). The strongest association was identified on chromosome 3B, where 60 significant markers (LOD > 3) were mapped, including 45 within the region between 11.9 Mbp to 28.5 Mbp, co-located with the previously defined *Wsm2* region (15.0 - 19.0 Mbp) (Table S6). In addition, two other significant markers were mapped to chromosome 3D at physical positions 4,397,505 bp and 5,446,355 bp. Sequence alignment showed that this region is not syntenic to the *Wsm2* locus on chromosome 3B (Figure S2). Five other significant markers were mapped to chromosome 7B, and one significant marker was mapped to chromosome 5B (Table S6). The results confirmed that ‘Snowmass’ contains the *Wsm2* variant and that this locus confers WSMV resistance. Therefore, ‘Snowmass’ can be used to identify and characterize genetic variation underlying the *Wsm2* locus.

**Figure 2.**
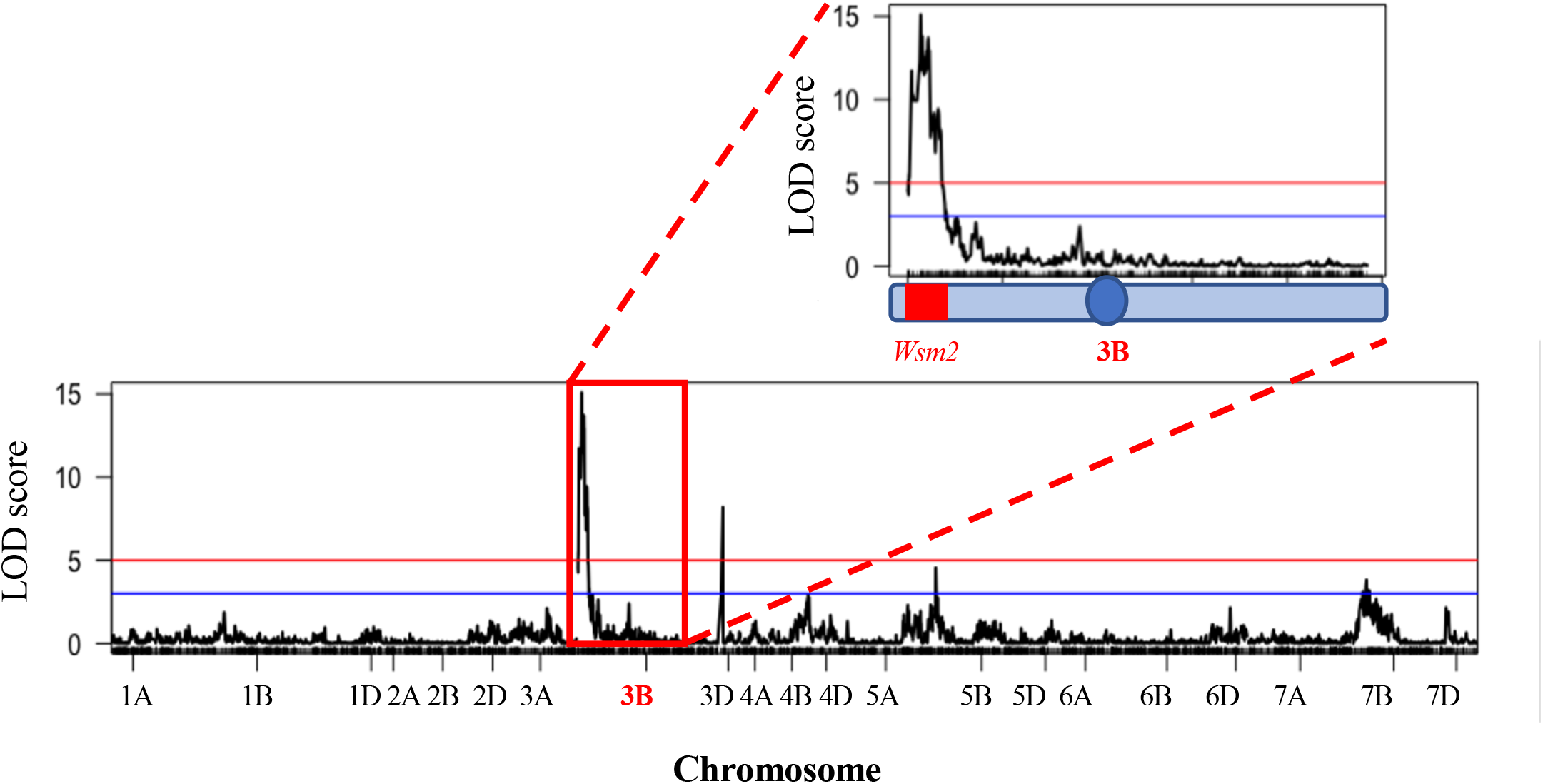
QTL mapping for WSMV resistance in a ‘Snowmass’ × ‘Antero’ doubled haploid population (n = 116). Horizontal blue line indicates LOD = 3 (*P* < 1e-3) and horizontal red line indicates LOD = 5 (*P* < 1e-5). Red box indicates chromosome 3B. The blue circle indicates the position of the chromosome 3B centromere and the *Wsm2* region is highlighted in red.

#### 3.1.3 Exome sequencing revealed genetic and copy number variation at the *Wsm2* locus

Exome sequencing reads from ‘Snowmass’ were mapped to the IWGSC RefSeq v1.0 genome to characterize natural genetic variation in protein-coding genes between ‘Chinese Spring’ (*Wsm2-*) and ‘Snowmass’ (*Wsm2+*). Within the *Wsm2* interval, 1,191 SNPs (96.0%) and 50 small Indels (4.0%) were identified (Table S7). This translates to a rate of one variant per 126 bp of coding sequence, higher than the mean rate of one variant per 503 bp across the whole exome, as well as in six other 4 Mbp regions sampled from different regions of the genome (Table S8). However, a high rate of variation in this region was also observed in six other wheat genotypes (‘Antero’, ‘Brawl’, ‘Byrd’, ‘Hatcher’, ‘CO940610’, and ‘Platte’), ranging from one variant per 125 bp to 189 bp exome length (Table S8). None of these varieties exhibit resistance to WSMV, suggesting that the high rate of variation is unrelated to the presence of *Wsm2*.

The variants within *Wsm2* are predicted to induce 4,382 genetic effects within spliced transcript sequences (Table S7). Six high-impact variants were predicted, leading to either premature introduction of a stop codon within the coding sequence or a shift in the open reading frame (Table S7). Of the six high impact variants in ‘Snowmass,’ four were also present in the WSMV susceptible parent ‘Antero’ (Table S7). One of the genetic variants unique to ‘Snowmass’ is a 2 bp insertion in *TraesCS3B02G042400LC*, a non-translating gene, and the other is a 10 bp deletion in *TraesCS3B02G038300* (Bowman-Birk trypsin inhibitor) that introduces a premature stop codon at amino acid 156 (T156*) and is likely to encode a non-functional protein (Table S7).

To search for structural variation within the *Wsm2* interval in ‘Snowmass’, exome sequencing read depth from ‘Snowmass’ was compared to the mean depth from the exomes of six other wheat varieties. Of the 142 candidate genes within *Wsm2*, ten exhibited copy number variation (Figure 3, Figure S3). Six genes underwent an expansion in ‘Snowmass’ and are predicted to have two to three copies compared to the mean coverage in the reference exome set (Figure 3). These include a cluster of four adjacent genes encoding UDP-glycosyltransferase proteins and a gene encoding an NBS-LRR type disease resistance protein (Figure 3).

**Figure 3.**
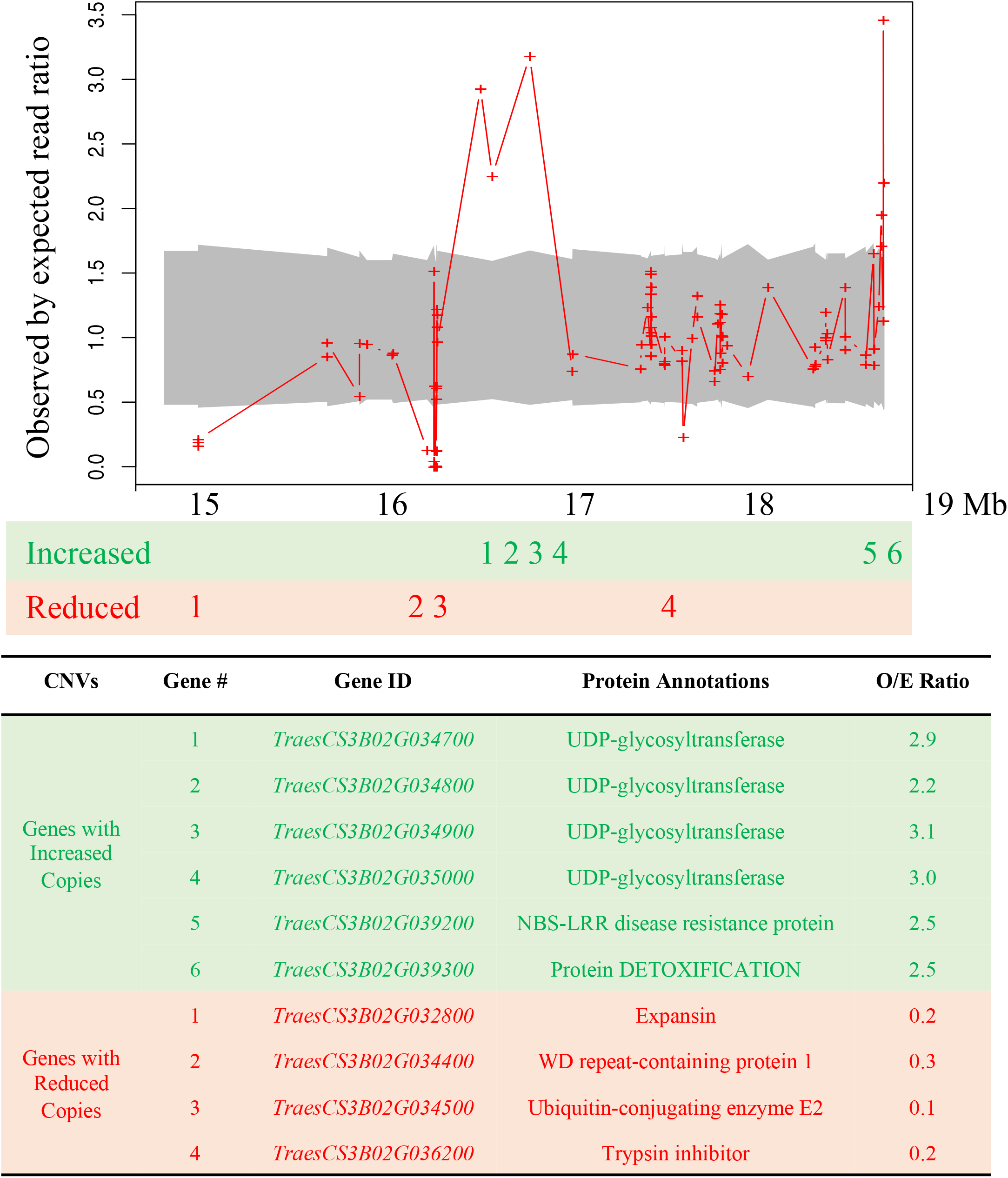
Copy number variation at the *Wsm2* locus in ‘Snowmass’. Values indicate the observed by expected read ratio between the ‘Snowmass’ exome and a reference set of exomes from six other wheat varieties. The 95% confidence interval is marked by grey shadow. Red crosses indicate a minimum number of ten reads mapped to this region. The labels below indicate the order of genes that are predicted to exhibit either increased or reduced copy number. The table describes the protein annotation and observed by expected read ratio (O/E ratio) averaged for all reads that mapped to each gene. Genes highlighted in green are predicted to have more copies in ‘Snowmass’, whereas genes highlighted in red are predicted to have fewer copies in ‘Snowmass’.

### 3.2 Transcriptomics to characterize host response to WSMV infection

#### 3.2.1 WSMV accumulation in *Wsm2+* and *Wsm2-* genotypes

To compare the accumulation of WSMV in resistant and susceptible wheat, RNA was extracted from whole leaf tissues from ‘Snowmass’ (*Wsm2*+, WSMV resistant) and ‘Antero’ (*Wsm2*-, WSMV susceptible) at four time points after WSMV inoculation (0, 5, 10, and 15 dpi, Figure 4A). WSMV accumulated in both genotypes throughout the time course, but at a much lower rate in *Wsm2*+ compared to *Wsm2*- (Figure 4A). There were no significant differences in WSMV coat protein transcript levels between genotypes at either 0 or 5 dpi (*P* > 0.05), but at both 10 dpi (4.4-fold, *P* < 0.001) and 15 dpi (4.7-fold, *P* < 0.05) *Wsm2*+ contained significantly lower levels of WSMV transcripts than *Wsm2*- (Figure 4A). This result was consistent with visual symptoms; whereas *Wsm2*- individual plants showed characteristic streaked, and mosaic patterns on their leaves beginning at 10 dpi, the leaves of *Wsm2*+ individuals remained asymptomatic throughout the time course (Figure 4B).

**Figure 4.**
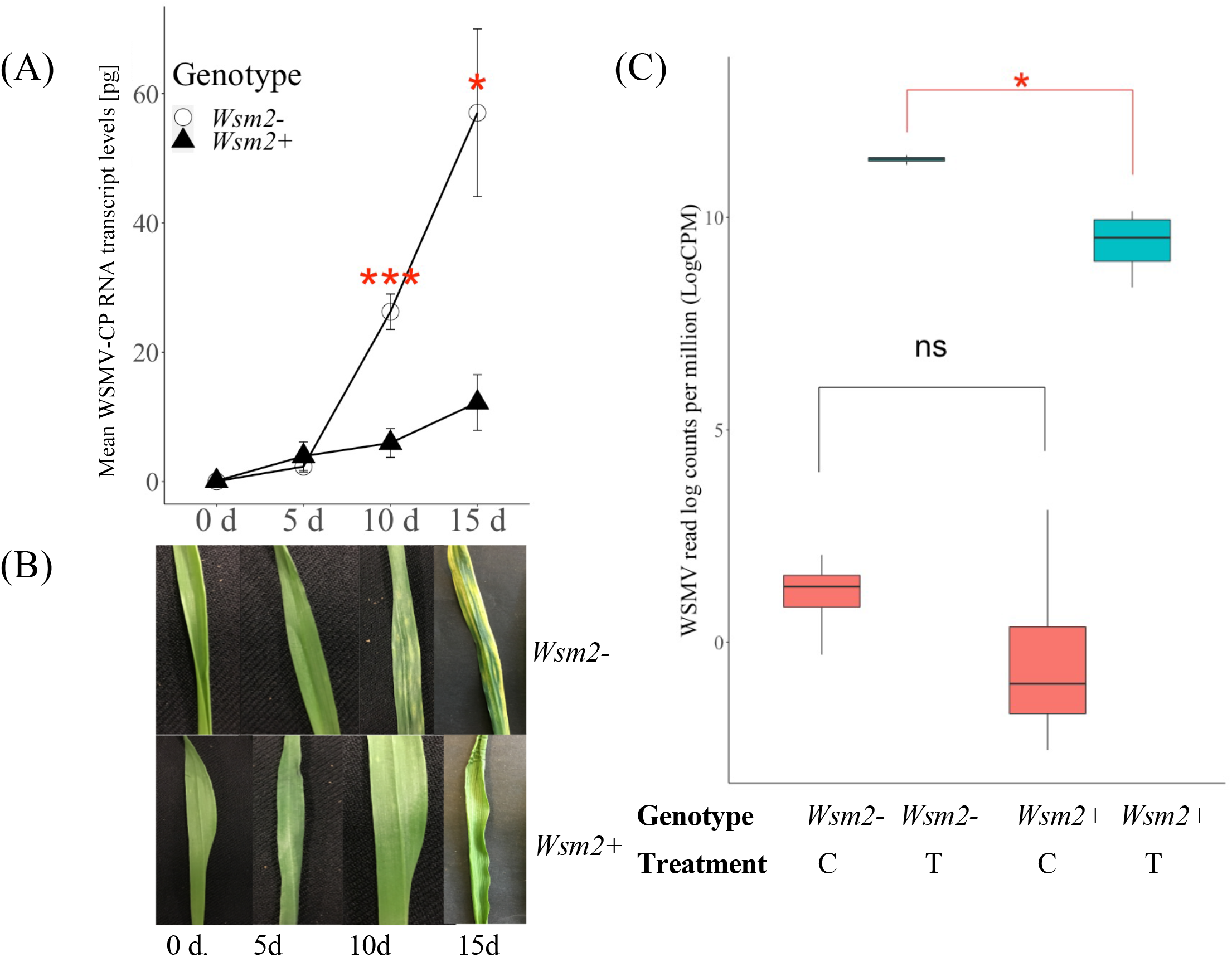
Characterization of the response of *Wsm2*+ and *Wsm2*- genotypes to WSMV infection. **A)** WSMV quantification in ‘Snowmass’ (*Wsm2*+) and ‘Antero’ (*Wsm2*-) leaf tissue before (0), 5-, 10- and 15- day post inoculation (dpi, n = 5). Error bar indicates standard error (SE, n = 5). **B)** Phenotype of leaves in ‘Snowmass’ and ‘Antero’ 0, 5, 10, and 15- dpi. **C)** Quantification of WSMV reads (log counts per million) in RNA-seq samples (n = 4) collected at 10 dpi from leaf tissue under four conditions: *Wsm2*- (C), *Wsm2-* (T), *Wsm2*+ (C) and *Wsm2*+ (T), T indicates WSMV-treated condition, C indicates mock-treated condition. Two tailed t-test were performed to compare between genotypes. * = *P* < 0.5, ** = *P* < 0.01, *** = *P* < 0.001, ns = not significant.

#### 3.2.2 Summary statistics of the RNA-seq experiment

Based on the time course results, 10 dpi was selected to characterize the early transcriptomic response of wheat plants to WSMV infection. In this RNA-seq study, 16 samples were collected from the leaf tissue of two wheat genotypes (*Wsm2*+, *Wsm2*-, Table S1) and under two treatments (WSMV-treated, mock-treated). After adaptor trimming and removal of low-quality reads, an average of 26.7 million reads were retained (Table S9) and mapped to a combined wheat-WSMV reference genome. Although WSMV reads were detected in all samples, the levels were much higher in WSMV-treated (T) samples (mean 10.4 LogCPM) compared to mock-treated (C) samples (mean 0.4 LogCPM, Table S10). WSMV levels were also significantly higher in *Wsm2*- (T) samples (mean 11.4 LogCPM) than in *Wsm2*+ (T) samples (mean 9.4 LogCPM, *P* < 0.05, Figure 4C), consistent with qRT-PCR results at 10 dpi (Figure 4A).

The overall mapping rate was 97.2% across 16 samples. The average unique mapping rate for *Wsm2*- wheat samples was 84.5 ± 3.3%, whereas for *Wsm2*+ samples the rate was 81.0 ± 1.3% (Table S9). In the principal component analysis (PCA), PC1 explained 45% of variance in the overall transcriptome between samples and most samples were separated according to treatment (Figure S4). There were three ambiguous samples, of which two belong to *Wsm2*+ (T) (R2 and R4) and one belongs to *Wsm2*- (C) (R4) (Figure S4). The counts of WSMV reads in the two ambiguous *Wsm2*+ (T) samples were 8.4 and 9.2 LogCPM compared to an average of 10.0 LogCPM in other *Wsm2*+ (T) samples (Table S10), suggesting that transcriptome variation in these samples may be due to variation in WSMV inoculation and infection. Samples were not grouped according to their genotype, indicating that transcriptomic variation between genotypes was comparatively smaller than between treatments.

#### 3.2.3 Host transcriptomic response to WSMV infection

To characterize host transcriptomic responses to WSMV infection, differentially expressed genes (DEGs) between mock- and WSMV-treated susceptible materials were analyzed (Table S11). In total, 8,975 DEGs were detected between *Wsm2*- (T) and *Wsm2*- (C) conditions (*P* < 0.01), of which 5,031 were up-regulated and 3,944 were down-regulated after WSMV infection (Figure 5A).

**Figure 5.**
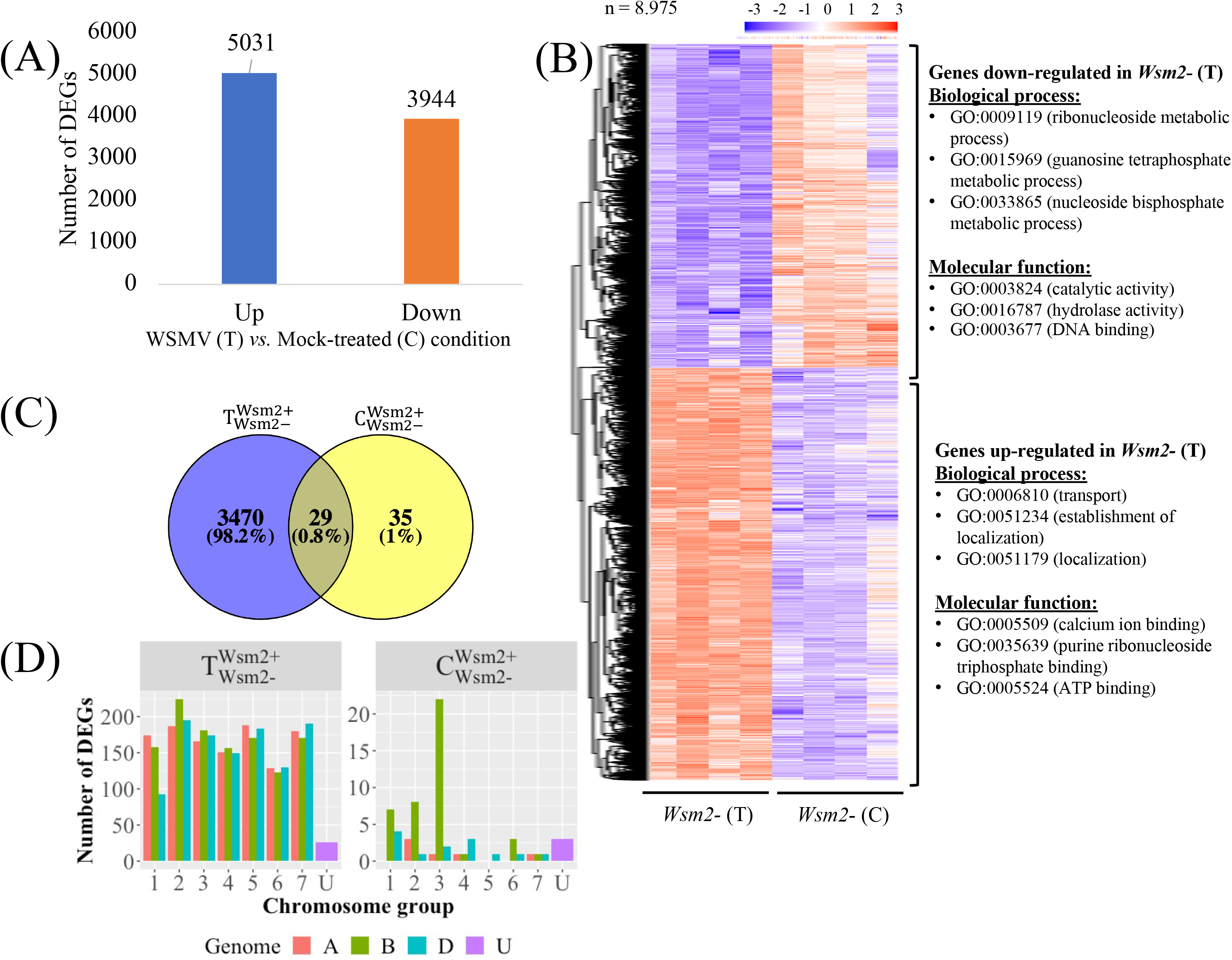
Overview of differentially expressed genes between mock (C) and WSMV (T) treatments, and between *Wsm2*+ and *Wsm2*- genotypes using IWGSC RefSeq v1.0 as a mapping reference. **A)** Number of up- and down-regulated DEGs (*P*adj < 0.01) between *Wsm2*- (C) *vs. Wsm2*- (T) samples. Up refers to genes that were more highly expressed in *Wsm2*- (T), whereas Down refers to genes that were more highly expressed in *Wsm2*- (C). **B)** Heatmap of 8,975 DEGs from comparison of *Wsm2*- (C) *vs. Wsm2*- (T) samples. The expression values are normalized by setting the mean of every row to zero and the standard deviation of every row to one. Hierarchical clustering separated these into DEGs that are either upregulated (n = 5,031) or downregulated (n = 3,944) in WSMV-treated conditions. The top three enriched GO terms (biological process and molecular function) for each row cluster are shown on the right. **C)** Venn diagram of total DEGs (*P*adj < 0.01) between genotypes 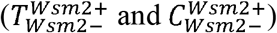. **D)** Number of DEGs comparing *Wsm2*+ (T) *vs. Wsm2*- (T) and *Wsm2*+ (C) *vs. Wsm2*- (C) conditions, based on each gene’s chromosomal location. A, B, D and Unknown genomes are color coded.

Down-regulated genes were most significantly enriched (*P* < 0.01) for biological process gene ontology (GO) terms relating to ‘photosynthesis’ (GO:0015979) and metabolic processes such as ‘purine ribonucleoside metabolic process’ and ‘lipid metabolic process’ (Figure 5B, Table S12), indicating that host plants suppress growth-related metabolic activity in response to WSMV infection. By contrast, up-regulated genes were significantly enriched for biological process GO terms related to ‘transport’ (GO:0006810) and ‘localization’ (GO:0051179) (Figure 5B, Table S12). Among these up-regulated genes were three *Pathogenesis-related 1* (*PR1*) genes, six *PR2* (β-1,3-glucanase) and *PR3* (chitinase) genes, four *PR5* (thaumatin-like protein) genes and fifteen *PR9* (peroxidase) genes, together with two homoeologous genes encoding RNA-binding proteins (*P* < 0.01, Table S13), suggesting their potential role in host response to viral infection.

#### 3.2.4 Difference in transcriptomic response between genotypes

To characterize transcriptional changes between genotypes, DEGs were analyzed under mock-treated 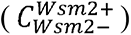 and WSMV-treated 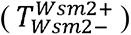 conditions. Only sixty-four genes were differentially expressed between genotypes in mock-treated conditions (Figure 5C), of which 28 were more highly expressed in *Wsm2+* genotypes, and 36 were more highly expressed in *Wsm2-* genotypes (Table S11). Twenty-two (34.4%) of these genes are located on chromosome 3B (Figure 5D, Table S14), and six are among the 142 candidate genes underlying *Wsm2* (See section 3.3).

In comparison, 3,499 genes were differentially expressed between genotypes under WSMV- treated conditions, of which 1,920 were more highly expressed in *Wsm2*+ genotypes, and 1,579 were more highly expressed in *Wsm2*- genotypes (Table S11). Twenty-nine genes were differentially expressed between genotypes in both mock- and WSMV-treated conditions, while 3,470 genes were differentially expressed only after WSMV-treated conditions (Figure 5C). These results indicate that the host plant response to WSMV infection varies depending on the presence or absence of *Wsm2*. These 3,470 DEG are significantly enriched for GO terms relating to different metabolic processes (GO:0046128, GO:0072521, GO:0033865) and catalytic activity (GO:0003824, GO:0016757) (Table S12), indicating that the differences between genotypes in the days following WSMV infection include modified cellular metabolism and catalytic activity. However, examination of enriched GO terms for these DEGs did not find any ‘defense response,’ ‘hormone regulation,’ ‘signaling transduction,’ ‘cell wall biogenesis’, or ‘photosynthesis’ related terms (Table S12).

### 3.3 Analysis of transcriptomes to identify candidate genes underlying *Wsm2*

#### 3.3.1 Examination of candidate genes within *Wsm2* found six genes differentially expressed between genotypes

Of the 142 annotated candidate genes within the *Wsm2* interval in the IWGSC RefSeq v1.0 genome assembly, six were differentially expressed between *Wsm2+* and *Wsm2-* genotypes in both WSMV-treated (T) and mock-treated (C) conditions (Figure 6A). The differential transcript levels at 10 dpi between genotypes of five of these genes were validated using qRT-PCR (Figure 6B), demonstrating the reliability of RNA-seq in quantifying transcript levels. Some were also significantly differentially expressed at earlier or later time points following inoculation, demonstrating they exhibit sustained differences in expression between genotypes (Figure 6B).

**Figure 6.**
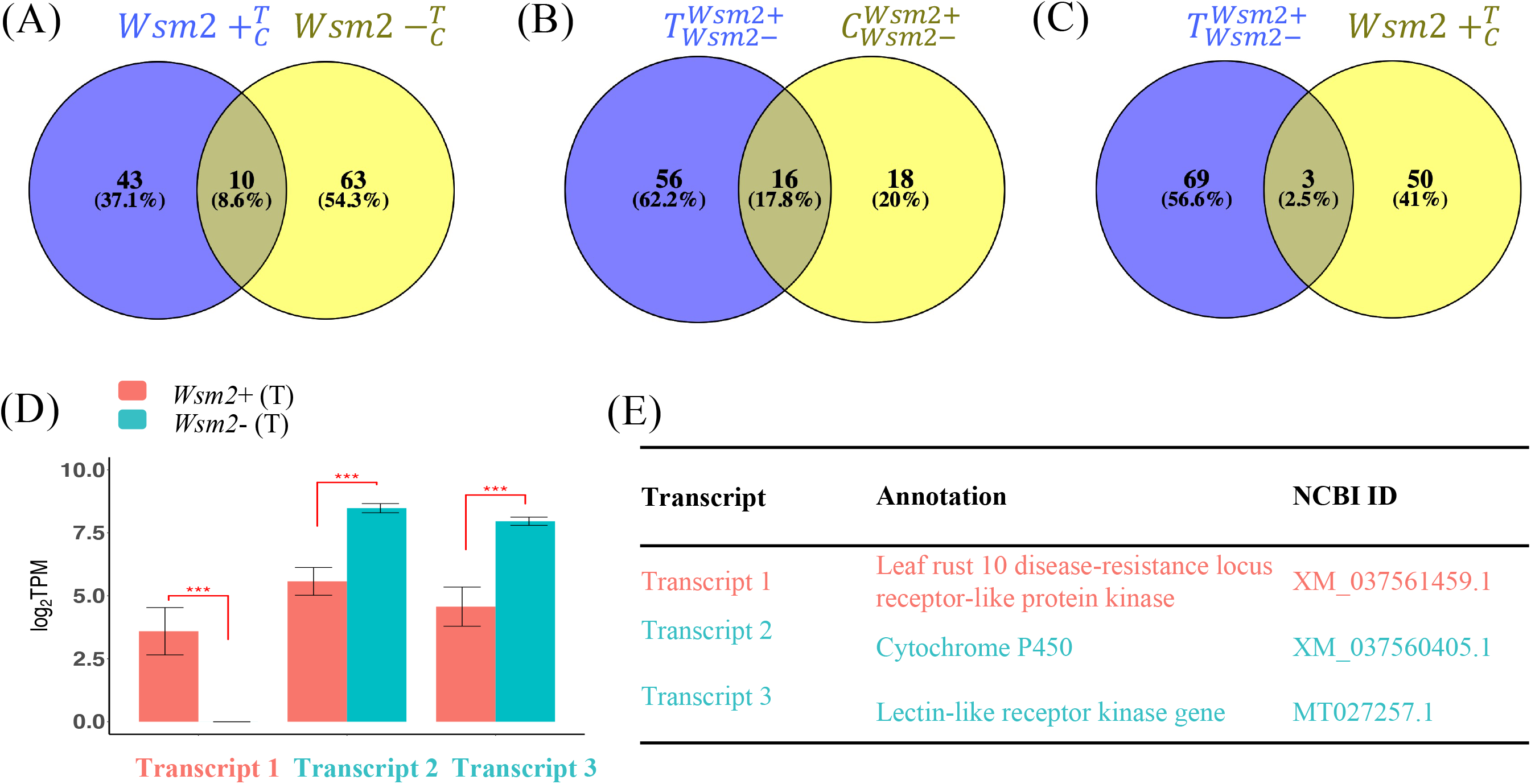
Transcript levels of six DEGs within the *Wsm2* region. **A)** log_2_TPM values of six DEGs at 10 dpi quantified by RNA-seq. Values were color-coded by their sample group. *Wsm2+* under WSMV-treated (*Wsm2+* T) and mock-treated condition (*Wsm2*+ C); *Wsm2*- under WSMV-treated (*Wsm2*- T) and mock-treated condition (*Wsm2-* C). *** = *P* < 0.001 **B)** Relative expression in fold-change *ACTIN* levels for five of the DEGs at four time points (0, 5, 10, and 15-dpi), quantified by qRT-PCR. * = *P* < 0.05. **C)** Annotation for the six DEGs.

Two genes, encoding a WD repeat-containing protein 1 (*TraesCS3B02G034400*) and Ubiquitin-conjugating enzyme E2 (*TraesCS3B02G034500*), were more highly expressed in *Wsm2-* (Figure 6C, Table S15). Both genes exhibit reduced copy number in ‘Snowmass’ (Figure 3), potentially explaining their lower transcript levels in *Wsm2*+ genotypes. The other four candidate genes were more highly expressed in *Wsm2*+ genotypes, and encode a putative receptor kinase (*TraesCS3B02G032400*), a SUF system FeS protein (*TraesCS3B02G035600*), NBS-LRR type protein with homology to RPM1 (*TraesCS3B02G035800*), and a Chaperone protein DnaK (*TraesCS3B02G035900*) (Figure 6C).

None of the four UDP-glycosyltransferase genes predicted to exhibit increased copy number in ‘Snowmass’ (Figure 3) were differentially expressed between genotypes (Table S15), suggesting these genes are unlikely to contribute to WSMV resistance. Despite their potential roles in biotic stress resistance, all twelve BBI genes within the *Wsm2* interval exhibit low expression levels (TPM < 0.4) across all samples and were not differentially expressed between genotypes (Table S15).

#### 3.3.2 *De novo* assembly of unmapped reads revealed transcripts absent from the wheat reference genome

To identify potential causative genes underlying *Wsm2* that are absent from the IWGSC RefSeq v1.0 genome, a *de novo* assembly of RNA-seq reads that did not map to this reference was performed. A total of 23,066,200 unmapped reads (5.4% of all reads, Table S9) were combined from all 16 samples and assembled into 161,210 non-redundant transcripts.

The unmapped RNA-seq reads from each sample were mapped back to the *de novo* assembled transcriptome, revealing 245 transcripts that were differentially expressed in at least one of the four pairwise comparisons (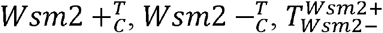, and 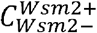, Table S16). Of these, 56 were annotated as sequences from non-plant species, including 42 matching WSMV, and were excluded from the analysis. Of the remaining 189 transcripts, nine were absent or expressed at very low levels in *Wsm2*- genotypes (defined as transcript levels < 0.2 TPM in both WSMV- and mock-treated conditions, Table S17). Using BLAST, these nine transcripts were confirmed to be absent from ‘Chinese Spring’ and, with two exceptions, absent from the genomes of ten other wheat varieties (Tables S16 and S17). Among these *Wsm2*+-specific transcripts are one that encodes an LRR receptor, one that encodes a BBI trypsin inhibitor, and three transcripts that encode Leaf rust 10 resistance proteins (Table S17). Sixteen other transcripts were absent from *Wsm2*+ genotypes, including one predicted to encode a negative regulator of resistance protein (Table S17).

There were 116 transcripts differentially expressed between WSMV-treated and mock-treated conditions, including 43 only in the *Wsm2+* genotype, 63 only in the *Wsm2-* genotype, and ten shared between both genotypes (Figure 7A). Additionally, a total of 90 transcripts were differentially expressed between genotypes (18 only in mock-treated conditions, 56 only in WSMV-treated conditions, and 16 in both conditions, Figure 7B). Among these transcripts, three were induced by *WSMV* treatment in the *Wsm2+* genotype (Figure 7C). One transcript encoding a leaf rust 10 disease resistance locus receptor-like protein kinase (XM_037561459.1) and was exclusively expressed in *Wsm2+* (T) conditions (TPM = 5, *P* < 0.001) (Figure 7D, Table S16). This transcript was present in six wheat varieties and in each case, was located on chromosome 3B, approximately 20 Mbp downstream of the left boundary marker for the *Wsm2* locus (Table S18).

**Figure 7.**
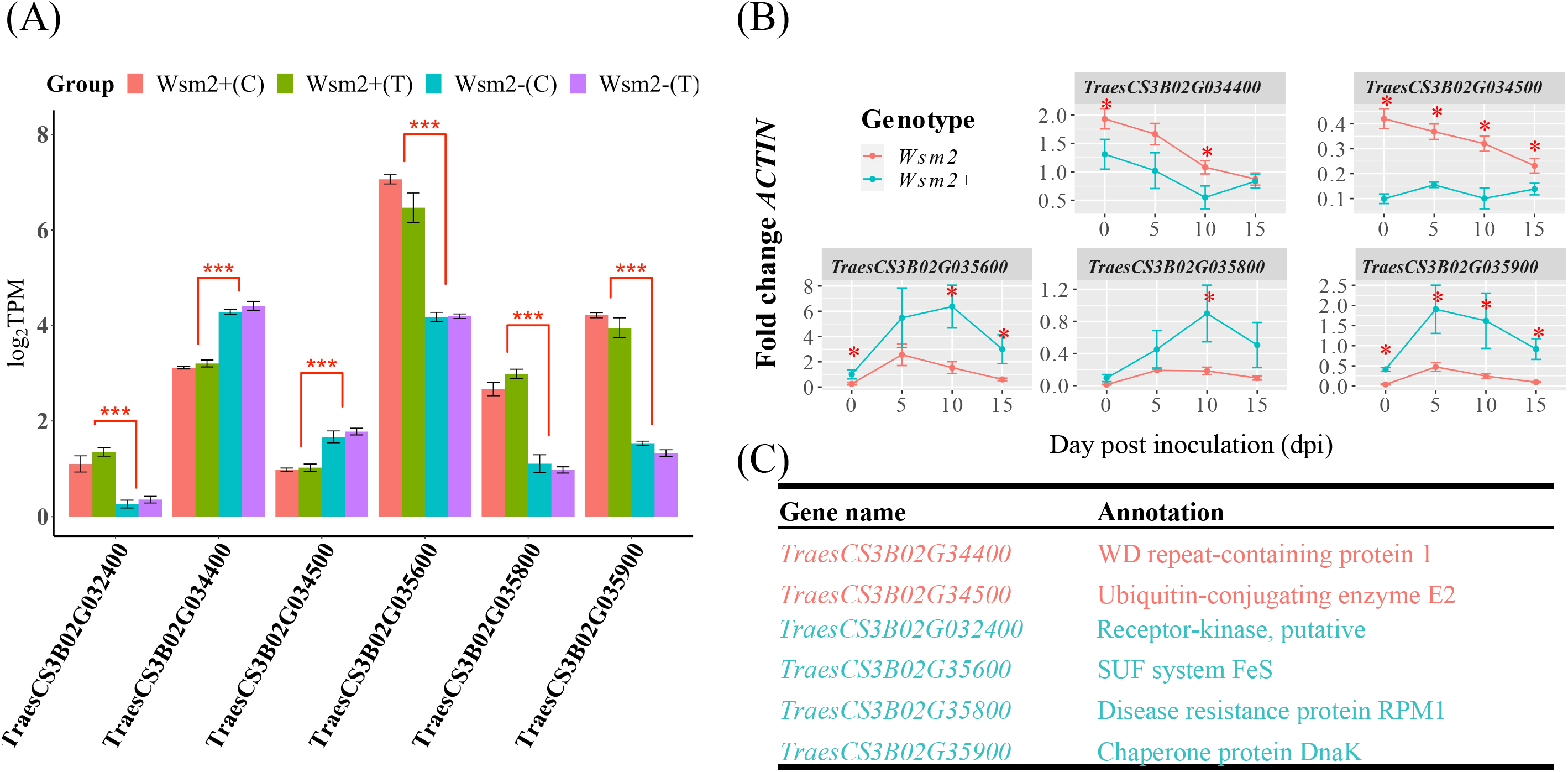
Expression analysis of *de novo* assembled transcripts absent from ‘Chinese Spring’. Venn diagrams show the total number of differentially expressed transcripts (*P*adj < 0.01) between **A)** 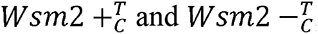, **B)** 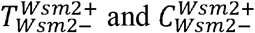, and **C)** 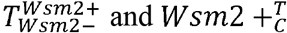. **D)** Expression in Log_2_TPM of three differentially expressed transcripts significant in both 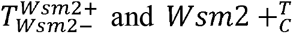 contrasts. *** = *P* < 0.001. **E)** Annotations of three differentially expressed transcripts.

The two other transcripts exhibited significantly higher expression in *Wsm2*- genotypes compared to *Wsm2+* genotypes in WSMV-treated conditions (*P* < 0.001), indicating they may be negatively associated with *Wsm2*-mediated resistance (Figure 7D, Table S16). One transcript encodes a cytochrome P450 (XM_037560405.1) and was present on the distal end of chromosome arm 3BL in ten wheat varieties with genome assemblies (Table S18). This is more than 800 Mbp from the *Wsm2* locus, suggesting it is unlikely to be located in the *Wsm2* region. The other transcript encodes a lectin-like receptor kinase gene (MT027257.1) and was absent from all wheat varieties, indicating that this likely represents a rare transcript in *Wsm2*+ genotypes (Table S18).

## 4 DISCUSSION

### 4.1 *Wsm2* lies in a highly dynamic region of the genome and is likely absent from many modern wheat varieties

The *Wsm2* locus was previously mapped to a 6.5 cM telomeric region of chromosome arm 3BS (Lu et al., 2012; Tan et al., 2017; Assanga et al., 2017), which corresponds to a 4.0 Mbp interval in the IWGSC RefSeq v1.0 wheat reference genome assembly (Table S4). The region of this genome is dynamic and variable, with at least eight haplotypic groups and several instances of large insertions and deletions among common wheat varieties (Figure 1). Telomeric regions of the chromosomes are associated with high recombination rates, resulting in frequent duplication and divergence events that potentially contribute to this variation (See et al., 2006; Saintenac et al., 2009). In addition to structural variation, exome regions of *Wsm2* in ‘Snowmass’ exhibited a higher variant rate than the average variant rate across all chromosomes (Table S8). Functionally constrained regions of the genome containing essential genes exhibit reduced mutation rates and are subject to stronger purifying selection 4/26/2022 2:33:00 AM. In contrast, resistance (*R*) genes tend to have a higher rate of variants than other genes, and many are located in clusters (Dolatabadian et al., 2017, 2020). It is more likely that the causative genes underlying *Wsm2* belong to an *R* gene type rather than an essential developmental role. The reduced purifying selection could contribute to the higher variant rate within this region.

The high rate of variants in the *Wsm2* region is shared between all studied varieties, including those that lack *Wsm2* (Table S8), and is consistent with *Wsm2* originating from a common wheat genetic background. However, based on their broad WSMV-susceptibility, it is likely that *Wsm2* is absent from all seventeen wheat genotypes with sequenced genomes, including IWGSC RefSeq v1.0, which is derived from ‘Chinese Spring’ (Table 1). This is in agreement with previous studies showing that ‘Chinese Spring’ is susceptible to WSMV (Tan et al., 2017) and that *Wsm2* is absent from wild *Brachypodium* accessions, a monocot ancestor of wheat (Zhang and Hua, 2018). Therefore, although the wheat pangenome is a powerful resource to exploit natural variation and characterize genetic variants associated with agronomic traits and stress resistance (Walkowiak et al., 2020), the absence of rare genetic variants from sequenced wheat germplasm might limit their application for some gene discovery projects.

Variation within the *Wsm2* region may explain previous findings of inconsistent marker order across this locus in different mapping populations (Tan et al., 2017) and may complicate the application of marker-assisted selection. Despite the diverse haplotypes in these seventeen wheat varieties, including four exhibiting the ‘AACT’ resistance haplotype, all genotypes exhibited a WSMV susceptible phenotype, although the result in ‘Jagger’ is ambiguous (Table 1). Taken together, the lack of association between these SNPs and WSMV resistance highlights the complexity of this locus and that marker-assisted selection should be approached with care. Cloning *Wsm2* would allow for the development of perfect markers to confirm its presence in different wheat germplasm for introgressing this resistance allele in breeding programs.

### 4.2 Candidate genes underlying *Wsm2*

That *Wsm2* lies in a dynamic region of the genome is consistent with a previous study showing multiple tandem duplication events involving clusters of BBI genes at this locus (Xie et al., 2021). Further genetic variation in the BBI family was detected in ‘Snowmass’. Compared to the wheat reference genome, ‘Snowmass’ is predicted to carry a non-functional allele of one BBI gene (*TraesCS3B02G038300*, Table S7) and a deletion of another (*TraesCS3B02G038300*, Figure 3). Although one *Wsm2*+ specific transcript annotated as a BBI was detected in ‘Snowmass’ (Table S17), overall, BBIs showed low transcript levels that were not were not significantly different between genotypes under WSMV treated conditions, suggesting they are unlikely to confer WSMV resistance.

*Wsm2* is a dominant allele (Lu et al., 2012), so it is likely that the variant is gain-of-function. Therefore, while the ‘Snowmass’ genome contains reduced copies of four genes within the *Wsm2* region (Figure 3), and lacks sixteen transcripts present in *Wsm2-* genotypes (Table S17), these are unlikely to be causative variants for *Wsm2*.

Six candidate genes within the *Wsm2* interval in ‘Chinese Spring’ were differentially expressed between genotypes (Figure 6). Two DEGs were up-regulated in *Wsm2+* throughout the WSMV infection timecourse (Figure 6B) and encode SUF system FeS (*TraesCS3B02G035600*) and Chaperone protein DnaK (*TraesCS3B02G035900*). The chaperone protein DnaK (HSP70) is known to respond to both biotic and abiotic stress by helping to prevent the accumulation of excessive newly synthesized proteins and ensure proper protein folding during their transition process (Park and Seo, 2015). Another up-regulated candidate in *Wsm2+* (*TraesCS3B02G035800*) encodes a CC-NB-LRR domain protein with homology to *RPM1*. In *Arabidopsis*, this protein recognizes the avirulence factor AvrRpm1 from the bacterial pathogen *Pseudomonas syringae* pv. *maculicola* 1 and triggers plant ETI defense responses (Grant et al., 1995).

It is also possible that *Wsm2* is a novel gene absent from the ‘Chinese Spring’ reference genome. Additional copies of tandemly duplicated UDP-glycosyltransferase genes were identified in ‘Snowmass’ (Figure 3). Although these additional copies were not associated with increased transcript levels 10 dpi infection (Table S15), they might be induced at other time points and play a role in plant defense against viral pathogens. UDP-glycosyltransferase genes have diverse roles in plant immunity against various types of pathogens. For example, UDP-glycosyltransferase proteins have been shown to function as negative regulators of the necrotrophic fungus *Botrytis cinerea* in *Arabidopsis* (Castillo et al., 2019), and to promote resistance to the hemi-biotrophic bacterial pathogen *Pseudomonas syringae* pv *tomato* carrying the *AvrRpm1* gene (Langlois-Meurinne et al., 2005). Moreover, the tomato gene *Twi1* gene, that encodes a glycosyltransferase, was shown to play a role in plant defense against tomato spotted wilt virus via secondary metabolites (Campos et al., 2019).

Through *de novo* assembly of unmapped RNA-seq reads, nine transcripts present in ‘Snowmass’ but absent from ‘Antero,’ ‘Chinese Spring,’ and most other wheat genomes were identified (Table S17). Five were annotated as disease-related genes (Leaf rust 10 disease resistance receptor protein kinase, Bowman Birk inhibitor, and LRR receptor-like kinase). To determine whether these genes are responsible for WSMV resistance, it will be necessary to map their position in the ‘Snowmass’ genome and perform functional characterization either by developing gene knockouts in ‘Snowmass’ or transforming each gene into WSMV-susceptible varieties. Three novel transcripts exhibited differential expression between genotypes and were also induced in response to WSMV infection and encode a leaf rust 10 disease resistance locus protein kinase, a lectin-like receptor kinase, and a cytochrome P450 (Figure 7, Table S18). Cytochrome P450 proteins function in phytoalexin biosynthesis, hormone metabolism regulation, and the biosynthesis of secondary metabolites and other defensive signaling molecules that regulate plant immunity against various pathogen types (Xu et al., 2015). The lectin-like receptor kinase gene (*LecRLK*) is a class of RLK that contains a lectin/lectin-like ectodomain that can bind to carbohydrates (Sun et al., 2020). *LecRLKs* are involved in plant basal defense against biotrophic and necrotrophic pathogens through carbohydrate signal perception, which triggers the PTI response (Sun et al., 2020). However, whether *LecRLKs* are also involved in ETI or play a role in plant response to viral infection remains unknown. The gene annotated as leaf rust 10 disease-resistance locus receptor-like protein kinase-like (*LRK10*) was first identified from wheat, providing resistance to the fungal pathogen *Puccinia triticina*, which causes wheat brown rust (Feuillet et al., 1997, 1998). The *LRK10* gene was later characterized as an NLR-class of *R* gene in wheat with a strong diversifying selected N-terminal CC domain, suggesting a complex molecular mechanism of pathogen detection and signal transduction (Loutre et al., 2009). Although no evidence suggests this *LRK10* is involved in plant defense response to viral pathogens, it is possible that this candidate could directly or indirectly interact with viral molecules and be involved in downstream signal transduction pathways important in immunity.

A more comprehensive analysis of genetic variation would require genomic sequencing of ‘Snowmass’. Although whole-genome sequencing would be feasible, targeted sequencing of a chromosome arm isolated by flow-sorting may be more cost-effective. This approach has been successfully applied to clone the broad-spectrum *Lr22a* leaf-rust resistance gene in wheat (Thind et al., 2017) and would be a valuable approach to studying the *Wsm2* locus and other rare alleles in wheat germplasm.

### 4.3 Insight into wheat host transcriptomic response to WSMV infection

Despite the threat that viral pathogens pose to crop production, we have only a limited understanding of host antiviral immune mechanisms in monocot crops (Mandadi and Scholthof, 2013; Huang, 2021). The current study revealed that metabolic processes and photosynthesis were suppressed in susceptible hosts following WSMV infection (Table S12). Our findings is consistent with previous studies, which showed that pathogen infection leads to the suppression of gene expression and protein production in photosynthetic processes (Bilgin et al., 2010; Göhre et al., 2012) due to the growth-to-defense tradeoff to optimize plant fitness and efficient use of resources (Huot et al., 2017).

*PR* genes are induced upon pathogen infection and encode proteins associated with host defense responses (Ren et al., 2020). *PR1* genes are considered markers for plant resistance to biotrophic pathogens (Van Loon and Van Strien, 1999) and three wheat *PR1* genes were induced by WSMV infection (Table S13), suggesting they may play a role in host response. The induction of *PR1* is usually associated with the accumulation of SA, a phytohormone involved in plant defense that may stimulate host antiviral responses through inhibition of viral replication, cell-to-cell movement, and long-distance movement (Singh et al., 2004). However, the role of SA in host responses to WSMV will require further validation. Members of the RNA-binding proteins (RBPs) family can also regulate SA-mediated plant immune responses in *Arabidopsis* (Qi et al., 2010). One wheat RBP protein (TaUBA2C) was recently shown to interact with cysteine-rich protein from the Chinese wheat mosaic virus, which activates downstream defense responses to inhibit viral infection (Li et al., 2022). In the current study, *TaUBA2C* was induced two-fold following WSMV treatment (Table S13), indicating that RBPs may play a role in the host response to WSMV infection.

In conclusion, genomic analyses indicate that *Wsm2* lies in a dynamic region of the wheat genome and is likely a rare allele in modern common varieties. This variation may complicate marker-assisted selection for virus resistance in breeding programs. Several candidate genes were identified, including genes found only in *Wsm2*+ genotypes. Sequencing the ‘Snowmass’ genome will facilitate the identification of *Wsm2*, which will expand our knowledge of viral resistance mechanisms in crops.

## 5 ACKNOWLEDGEMENTS

We are grateful to to Dr. Eduard Akhunov for providing exome sequencing data.

## 6 AUTHOR CONTRIBUTIONS

YX performed the experiment and bioinformatic data analysis and wrote the first draft of the manuscript. YX and SP designed the study and prepared the manuscript. PN and VN provided comments in the paper revision.

## 8 DATA AVAILABILITY STATEMENT

The raw sequence of RNA-seq reads is available from the NCBI Gene Expression Omnibus under accession number GSE190382. Full details of all data analysis steps and outputs are provided in supplementary files.

## 9 ETHICS APPROVAL AND CONSENT TO PARTICIPATE

Not applicable.

## 10 CONSENT FOR PUBLICATION

Not applicable.

---

**Figure S1.** Dot plot for pairwise alignment of genomic sequences for *Wsm2* in ‘ArinaLrFor’ (19.4 Mb – 23.4 Mb) and ‘Jagger’ (20.0 Mb – 24.0 Mb) *versus* IWGSC RefSeq v1.0 (15.0 – 19.0 Mb). Red dashed lines indicate the approximate position of the five SNP markers based on their location in the IWGSC Refseq v1.0 genome. Since SNP3 and SNP4 were close, their dashed lines were merged.

**Figure S2.** Dot plot for pairwise alignment of genomic sequences between significant QTL markers on chromosome 3D (4.0 – 6.0 Mb) and the *Wsm2* region on chromosome 3B (15.0 – 19.0 Mb) using IWGSC RefSeq v1.0 sequence.

**Figure S3**. High-resolution visualization of CNVs for candidate genes within the *Wsm2* region using ExomeDepth R package. **A)** Regions containing four candidate genes predicted to have reduced copies in ‘Snowmass’. **B)** Regions containing six candidate genes predicted to have increased copies in ‘Snowmass’.

**Figure S4.** PCA plot for 16 samples in the RNA-seq experiment. Genotype is indicated by shape, circle means *Wsm2*+ whereas triangle means *Wsm2*-; treatment type is indicated by color, red means WSMV-treated, blue means mock-treated. Each genotype and treatment combination have four biological replicates.

