## Supplemental_materials_1 for "Genomic And Molecular Characterization Of The *Wheat Streak Mosaic Virus Resistance Locus 2* (*Wsm2*) In Common Wheat (*Triticum Aestivum*. L)"

Supplementary Material

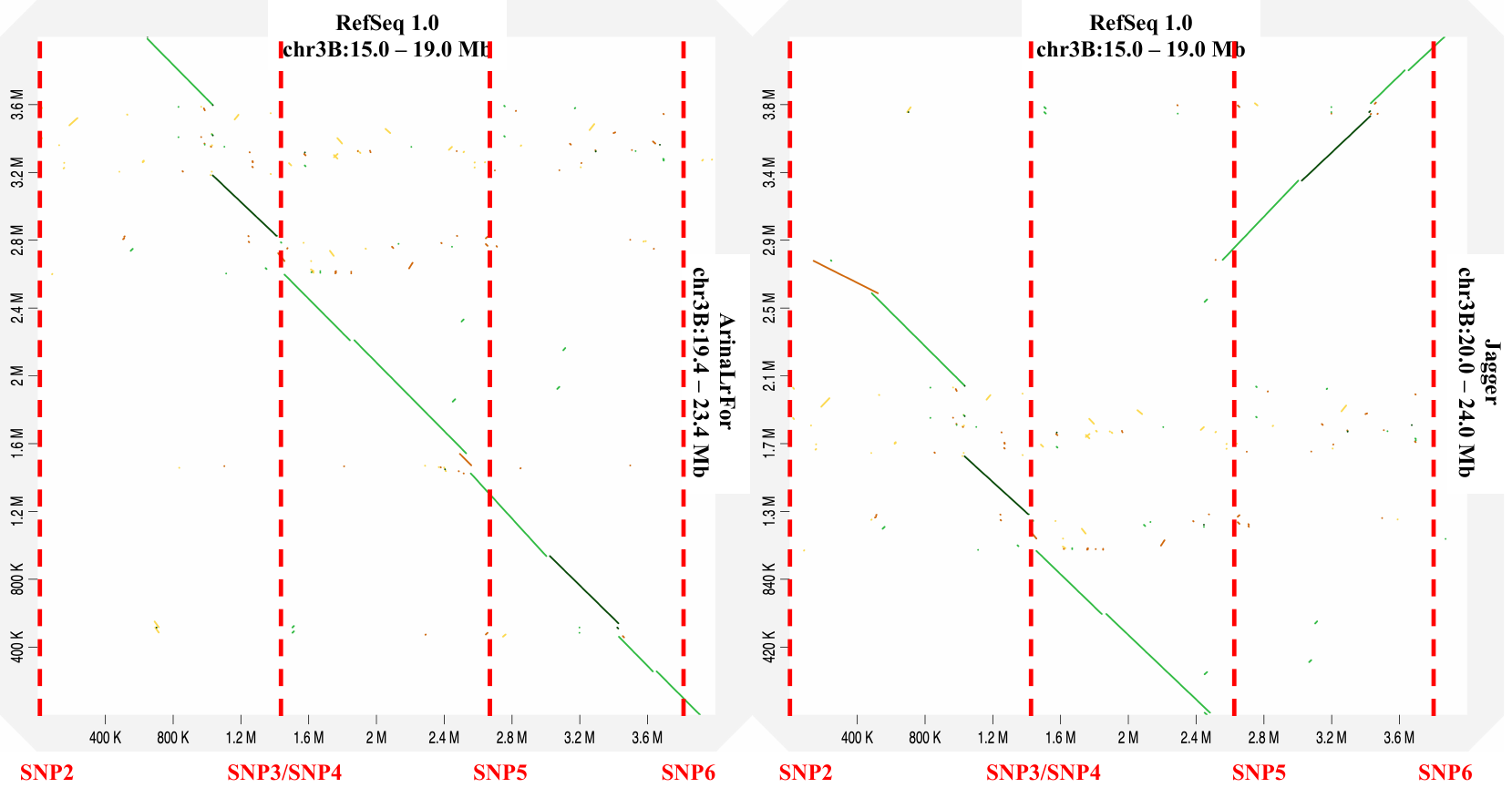

**Figure S1**. Dot plot for pairwise alignment of genomic sequences for *Wsm2* in ‘ArinaLrFor’ (19.4 Mb – 23.4 Mb) and ‘Jagger’ (20.0 Mb – 24.0 Mb) *versus* IWGSC RefSeq v1.0 (15.0 – 19.0 Mb). Red dashed lines indicate the approximate position of the five SNP markers based on their location in the IWGSC Refseq v1.0 genome. Since SNP3 and SNP4 were close, their dashed lines were merged.

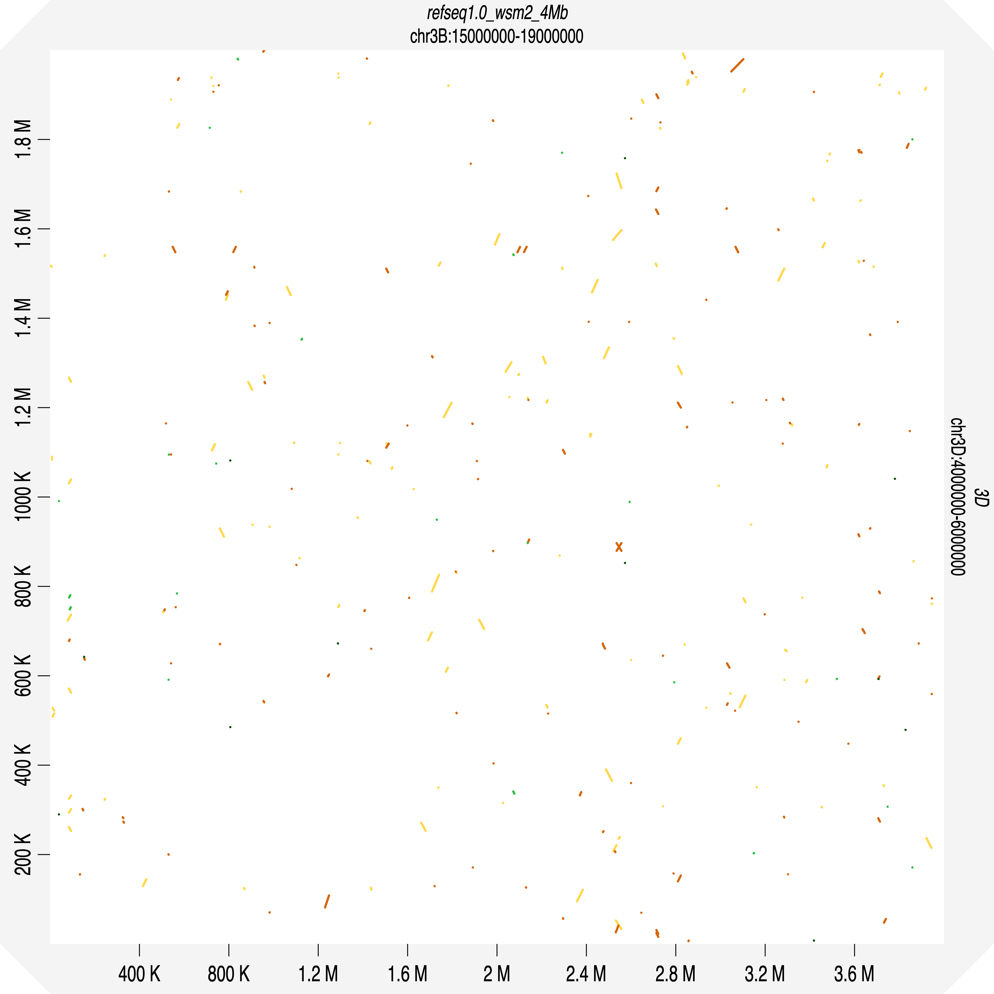

**Figure S2.** Dot plot for pairwise alignment of genomic sequences between significant QTL markers on chromosome 3D (4.0 – 6.0 Mb) and the *Wsm2* region on chromosome 3B (15.0 – 19.0 Mb) using IWGSC RefSeq v1.0 sequence.

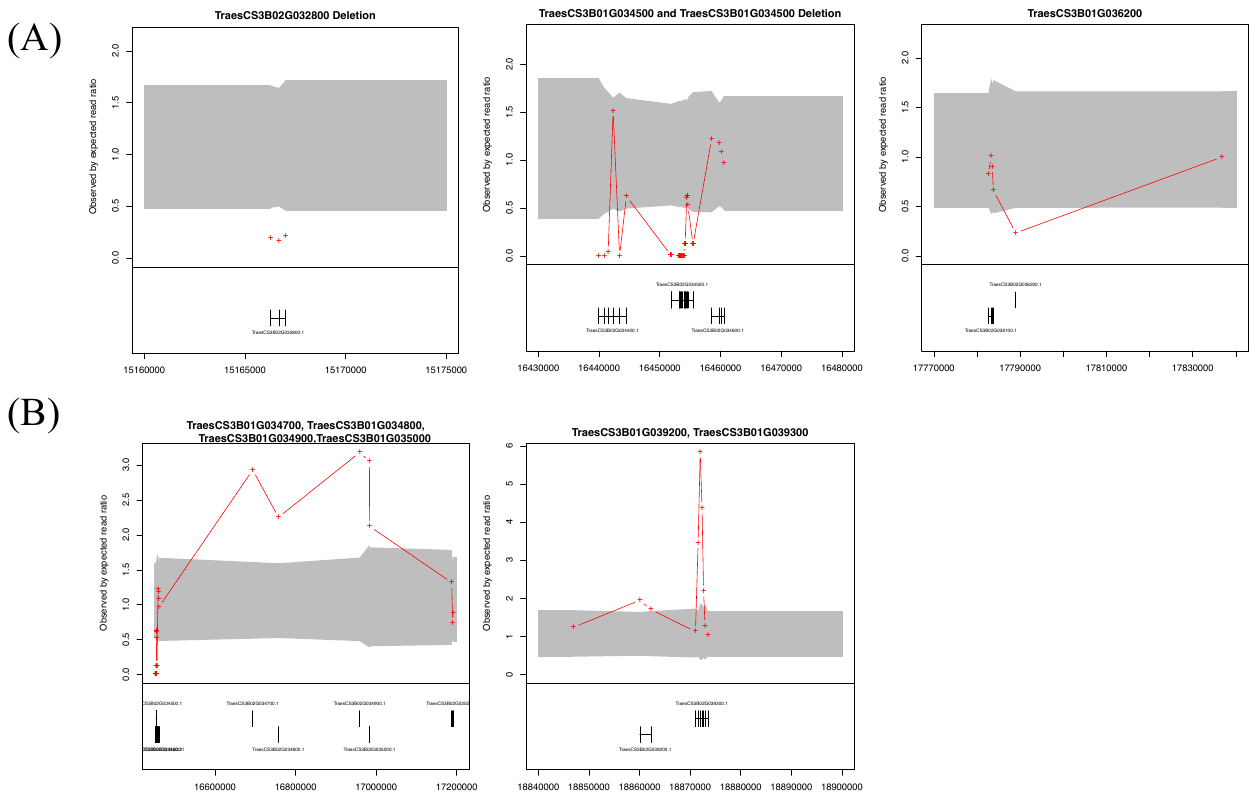

**Figure S3.** High-resolution visualization of CNVs for candidate genes within the *Wsm2* region using ExomeDepth R package. **A)** Regions containing four candidate genes predicted to have reduced copies in ‘Snowmass’. **B)** Regions containing six candidate genes predicted to have increased copies in ‘Snowmass’.

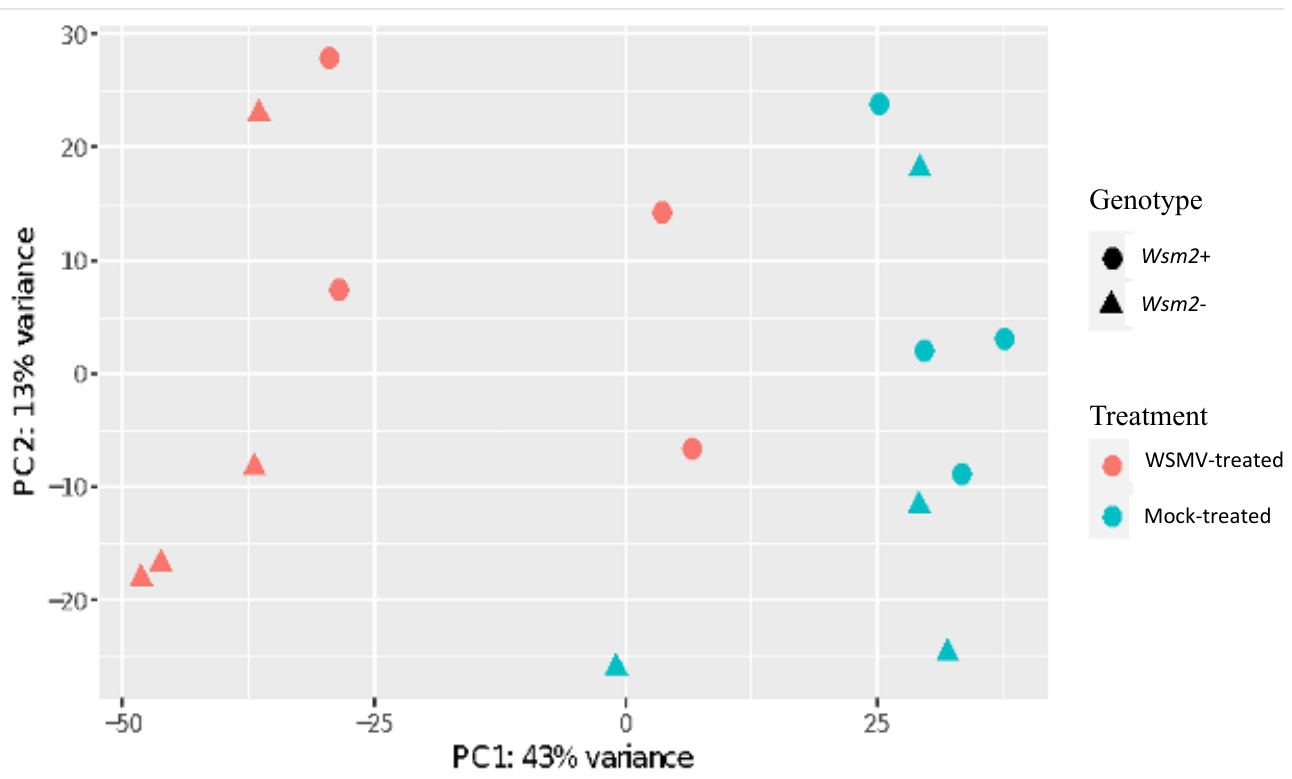
**Figure S4.** PCA plot for 16 samples in the RNA-seq experiment. Genotype is indicated by shape, circle means *Wsm2*+ whereas triangle means *Wsm2*-; treatment type is indicated by color, red means WSMV-treated, blue means mock-treated. Each genotype and treatment combination have four biological replicates.

**Table S1**. Haplotype and phenotype of selected individuals from the ‘Snowmass’ × ‘Antero’ doubled haploid population used for the RNA-seq study.

**Table S2.** Information for *Wsm2* flanking markers and three *Wsm2*-associated KASP markers (SNP1-SNP6).

**Table S3**. Primer and probe information for WSMV quantification and qRT-PCR of candidate genes.

**Table S4.** Information for 142 candidate genes within *Wsm2* interval, including their physical position in the IWGSC RefSeq v1.0 and IWGSC RefSeq v2.1 genome assemblies, gene ID and predicted functional annotations.

**Table S5**. Physical position of five *Wsm2* associated markers (SNP2-SNP6) in seventeen wheat varieties.

**Table S6**. Name and location of significant GBS markers (LOD > 3) associated with WSMV resistance from the linkage mapping in the Snowmass × Antero DH population.

**Table S7**. Polymorphisms and genetic variant effect for candidate genes within the *Wsm2* locus in ‘Snowmass’ determined from exome reads mapped to IWGSC RefSeq v1.0. Variants are separated by their type, region of the affected gene, impact and functional class. Full details of six high-impact polymorphisms, including two that are present in ‘Snowmass’ but absent from ‘Antero’, that are highlighted in red.

**Table S8**. Variant rate analysis (base pair of exome per variant) for each chromosome in ‘Snowmass’, as well as six selected 4 Mbp regions elsewhere in the genome. The variant rate for the *Wsm2* region (SNP2 to SNP6) in six other wheat varieties (‘Antero’, ‘Brawl’, ‘Byrd’, ‘CO940610’, ‘Hatcher’ and ‘Platte’) were also included.

**Table S9**. RNA-seq read numbers and mapping statistics for each sample mapped to the combined IWGSC RefSeq v1.0 and WSMV genome assembly and to the IWGSC RefSeq v1.0 genome assembly alone.

**Table S10**. Raw and normalized WSMV read counts (CPM and logCPM) for each of the 16 RNA-seq samples.

**Table S11**. Differentially expressed genes in four contrasts between treatment and genotype using IWGSC RefSeq v1.0 as a mapping reference. Mean transcript levels (TPM) for each condition, fold-change and FDR adjusted *P* values for each gene are provided.

**Table S12**. List of significantly enriched Molecular Function (MF) and Biological Process (BP) GO terms (*P* < 0.01) for up- and down-regulated DEGs between WMSV-treated *vs.* mock-treated samples and for the 3,470 unique DEGs between *Wsm2+* and *Wsm2-* genotypes in WSMV-treated conditions.

**Table S13**. Expression profiles of *PR* and defense related genes (in TPM) in wheat under four conditions (WSMV-treated *vs.* Mock-treated in two genotypes). The pairwise comparisons (fold change, log_2_ fold change, and *P*adj) were performed between WSMV *vs.* Mock treatment in *Wsm2-* samples, fold change > 1 and *Padj* < 0.01 were defined as up-regulated (Up) genes, whereas fold change < 1 and *Padj* < 0.01 were defined as down-regulated (Down) genes. Functional annotation, GO ID, and GO term for *PR1, PR2, PR3, PR5, PR9* were from Ren et al., (2020). The IWGSC gene ID for *SAPK7* and *TaUBPA2C* were from Li et al., (2022).

**Table S14**. Genomic position and functional annotation of the 22 DEGs on chromosome 3B that are significantly different in the $C_{Wsm2-}^{Wsm2+}$ comparison. DEGs within the *Wsm2* interval are highlighted in red.

**Table S15**. Gene expression and pairwise comparisons (TPM, log_2_ fold change, and *P*adj) for the 142 candidate genes underlying *Wsm2*. Genes that have increased copies in ‘Snowmass’ are in green text, and genes that have reduced copy number in ‘Snowmass’ are in red text. The Bowman-birk inhibitors characterized in Xie et al., 2021 are highlighted in yellow.

**Table S16**. Gene expression and pairwise comparisons (TPM, log_2_ fold change, and *P*adj) for *de novo* assembled transcripts from unmapped reads that were differentially expressed in any of the four pairwise comparisons. *P*adj < 0.01 are highlighted in pink, and the three differentially expressed transcripts that overlapped between $T_{Wsm2-}^{Wsm2+}$ and ${Wsm2+}_{C}^{T}$ are highlighted in red text.

**Table S17**. Information for differentially expressed transcripts (DETs) from plants that were absent or expressed at low levels (< 0.2 TPM) in one genotype. The DETs absent from one genotype were highlighted in yellow, *P*adj < 0.01 were highlighted in pink. Presence/absence analysis were performed for these transcripts in ten other wheat varieties. A dash (-) indicates that the transcript was absent, while the corresponding gene ID is provided if the transcript was present.

**Table S18**. Presence and absence analysis against ten wheat varieties for the three transcripts identified from *de novo* assembly of unmapped transcriptomes that exhibited differential expression between genotypes and were also induced in response to WSMV infection.
